# Deep Learning Reaction Network: a machine learning framework for modeling time resolved data

**DOI:** 10.1101/2024.07.31.606055

**Authors:** Nicolò Alagna, Brigitta Dúzs, Heinz Köppl, Andreas Walther, Susanne Gerber

## Abstract

Model-based analysis is essential for extracting information about chemical reaction kinetics in full detail from time-resolved data sets. Such analysis combines experimental hypotheses of the process with mathematical models related to the system’s physical mechanisms. This combination can provide a concise description of complex system dynamics and extrapolate kinetic model parameters, such as kinetic pathways, time constants, and species amplitudes. However, the process leading to the final kinetic model requires several intermediate steps in which different assumptions and models are tested, even using different experimental data sets. This approach requires considerable experience in modeling and data comprehension, as poor decisions at any stage of time-resolved data analysis (such as time-resolved spectra and agarose gel electrophoresis) can lead to an incorrect or incomplete kinetic model, resulting in inaccurate model parameters and amplitudes. The Deep Learning Reaction Network (DLRN) can rapidly provide a kinetic reaction network, time constants, and amplitude for the system, with comparable performance and, in part, even better than a classical fitting analysis. Additionally, DLRN works in scenarios in which the initial state is a non-emitting dark state and for multiple timescales. The utility of DLRN is also shown for more than one 2D system, as it performed well for both spectral and time-resolved agarose gel electrophoresis data.

## Main

Time-resolved techniques, such as spectroscopy or microscopy, are powerful and widely used tools for studying and investigating the kinetics of chemical reactions in complex systems in photobiology, photochemistry, and photophysics.^1–5^ These techniques generate one-, two-, or three-dimensional data sets, in which changes in the signal intensity are tracked as a function of time, which is one of the independent experimental variables. For example, time-resolved spectroscopy measurements are composed of two independent experimental variables, one being related to wavelength, wavenumber, magnetic field strength, and so forth. The other is the probing-time after excitation, which is used to track changes in the signal during the measurement. To reveal the processes underlying the observable changes in signal, computational and mathematical methods are necessary to analyze the network of chemical reactions and extrapolate the molecular properties linked to the system’s evolution^5,6^. The necessity of introducing these techniques arises from the fact that short-lived intermediate species can be difficult to detect with experimental measurements^7,8^. Indeed, many cases require different experimental techniques to be combined in order to identify specific reaction pathways. Furthermore, the chemical, biochemical, and cellular processes of complex systems require theoretical modeling to effectively disentangle information regarding their dynamics^9–14^. In practice, working with mathematical models for kinetic studies is almost mandatory, and the strategies used for data analysis can be classified into two families: model-independent and model-dependent^5,15–18^. Model-independent analysis is usually a global fitting technique (or global analysis, GA), which consists of a simultaneous analysis of multiple kinetic traces at different wavelengths using exponential functions. This method provides a functional description of a system’s kinetic profile, extrapolating the decay-associated amplitudes, primarily known in spectroscopy as decay-associated spectra (DAS), and the minimum number of time constants involved in the mechanism. Although GA is usually a necessary step in studying time-resolved systems, it is only an initial step in the analysis because it provides only partial information about the dynamics. The combination of a GA with the photochemical model is usually called global target analysis (GTA)^5,15,19^, which falls into the model-dependent category. This analysis is essential because kinetic modeling can quantitatively extrapolate kinetic mechanisms from time-resolved data, providing kinetic parameters such as species-associated amplitude (or species-associated spectra, SAS) and decay-time constants to describe complex photochemical dynamics.

Although GTA is a well-established method in which each species is connected to another via differential equations, this analysis is not always straightforward since it involves devising and testing kinetic reaction models based on previous knowledge and selected assumptions^5,15,18,20–22^. This can lead to an increase in the minimum number of parameters previously found by GA and generation of a set of models of which the most reasonable must be determined. This points to the challenge of using GTA to identify the correct kinetic model by the interplay between data and modeling approaches. Such problems scale increasingly poorly if the system becomes complex and has many variables to compute. For example, ATP-driven DNA dynamics^23–26^ show multi-step strand formation controlled by several enzymatic reactions. The kinetic model is challenging to determine, and the analysis requires the accurate selection and combination of chemical engineering and modeling approaches.

In recent years, the scientific community has increasingly worked on finding ways to study and describe chemical reaction networks (CRNs) using automated computational methods^27–30^. In particular, the application of machine learning (ML) methods to chemical reactions has significantly increased in popularity^6,28,31–35^. In this growing field of research, neural network models are trained in synthetic data and with already highly validated data to set the ground truth of the system such that the neural network learns to extrapolate complex patterns in the data through supervised learning. This capability of supervised neural networks to extract complex information from data is validated by comparing the network’s predictions, which are the outputs of the ML model after data processing, with expected results derived from unseen data with a known ground truth, making it possible both to evaluate the performance and accuracy of the network and its ability to analyze the system correctly. These methods are not only used for analyzing chemical reactions of known and new chemical compounds^36–40^, but have also been applied to the study of proteins^41–44^ to probe the free-energy and protein conformational landscape for extrapolating key features of biological systems.

This work presents DLRN (Deep Learning Reaction Network), a deep neural network based on an Inception-Resnet^45^ architecture created to combine GTA and ML methods. DLRN can disentangle all the kinetic information from a 2D time-resolved data set, giving the most probable kinetic model, related time constants for each pathway, and a maximum of four SAS for one timescale. Moreover, DLRN can analyze systems in which the initial state is a non-emitting dark state. DLRN was also tested to analyze multi-timescale 2D data, resulting in a good performance and the ability to identify multiple species for each timescale. DLRN performed well in the analysis of both time-resolved spectra and agarose gel electropherograms.

## Results

### DLRN analysis of synthetic time-resolved spectral data

Fig. 1 shows the entire DLRN pipeline for analyzing a time-resolved data set. The data set (Fig. 1a), representing a 2D image containing wavelength and time information, is sent through the neural network and analyzed using the model block computational pathway. This block analyzes the 2D signal, giving a one-hot encoding representation of the most probable model that DLRN can predict from 02 different models (**Error! Reference source not found.**b). Next, the one-hot encoding prediction is converted into the model matrix (**Error! Reference source not found.**c). An explanation of this representation of the kinetic model is given in the Supplementary Information (section 3). The model matrix, which provides information about the possible rate constants and pathways involved, is transformed into a binary matrix (Supplementary Information, section 3), where each *k*_i_ is substituted by a token of value 1, but without changing the signs in front of the rate constants. This passage is essential as the model prediction should operate for both time constants and amplitude analysis. The binary matrix is then sent as a second input for the DLRN time and amplitude blocks, which can extrapolate the numbers and values for the corresponding time constants (Fig. 1d) and amplitudes (Fig. 1e) linked to the kinetic model of the data set analyzed. The DLRN can extrapolate up to seven tau values for the time constants and a maximum of four spectra for the amplitude. The efficiency of the DLRN is illustrated by a comparison between the expected and predicted kinetic models (Fig. 1f), which shows the chemical reaction network and time constant values for each decay pathway in both cases. DLRN can predict the correct kinetic model with high-probability confidence (Fig. 1b) and, with good approximation, accurate time constants for each decay path.

**Fig. 1.**
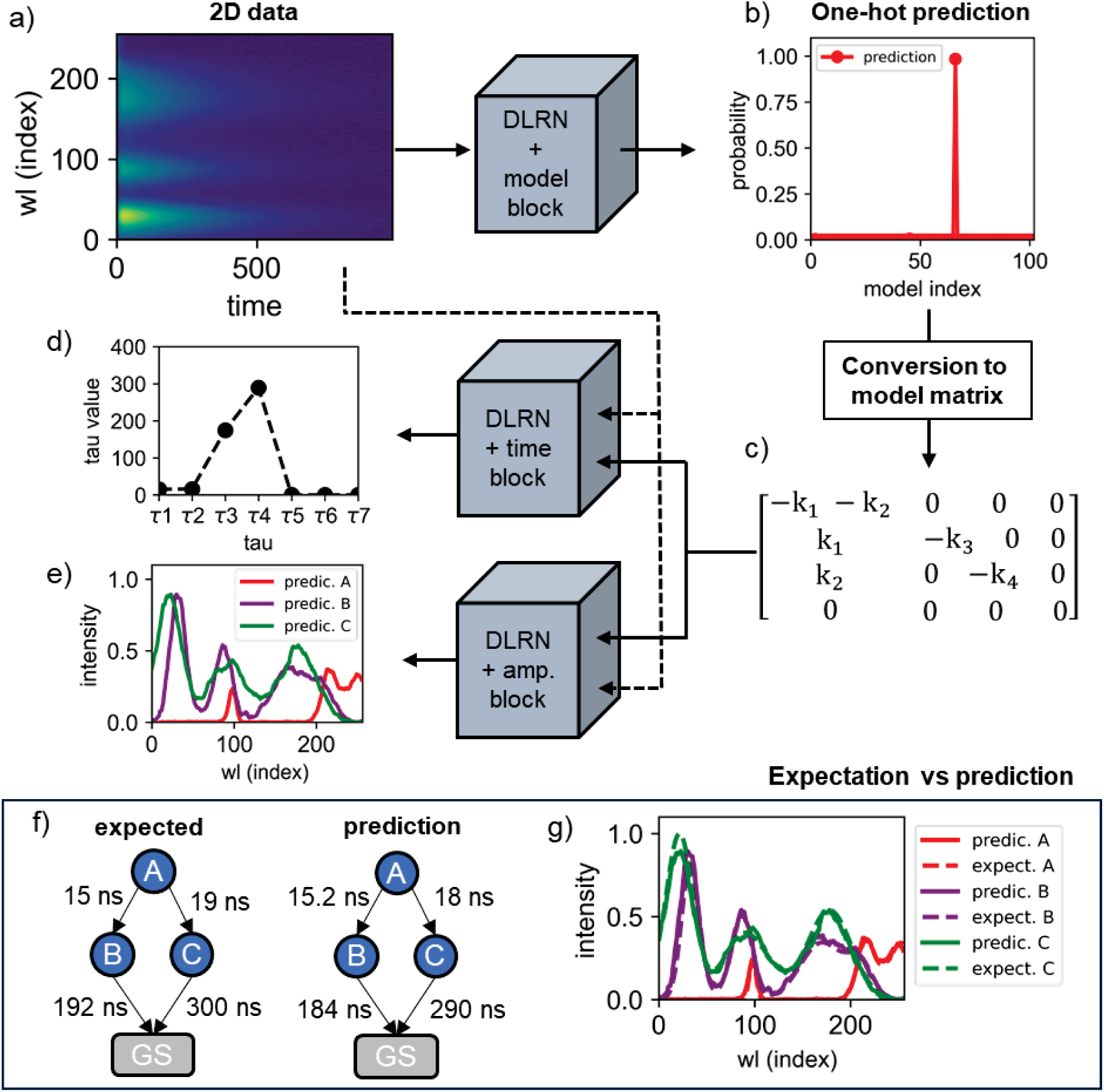
DLRN work-flow analysis of 2D images. **a,** Example of 2D spectra data used for the analysis. Wavelength (wl) are shown at the index position. **b,** One-hot encoding prediction obtained from DLRN using the model block. Each position represents a specific kinetic model. **c,** Conversion of the one-hot encoding prediction into a model matrix. This model matrix is used in both time and amplitude blocks for predicting time constants and amplitudes. **d,** Time constant predictions obtained from DLRN using the time block. **e,** Normalized amplitude prediction obtained from DLRN using the amplitude block. **f,** Comparison between expected and predicted models with the corresponding time constant for each decay pathway. **g,** Comparison between expected and predicted spectra. The predictions of model, time constants, and amplitudes match well the expected results.

Furthermore, the expected and DLRN-predicted amplitudes were compared (Fig. 1g). Again, the predictions show a good agreement between measured and simulated values.

Further comparisons between DLRN predictions and expected values are given in the Supplementary Information (section 2). To quantify the network’s performance in a larger number of samples, DLRN was tested on an evaluation batch containing 100,000 2D data sets. Table 1 shows the accuracy values obtained for the model, time constants, and amplitude predictions. The area metric *A*_m_ (see Methods) was used to quantify the accuracy of the regression analysis on the amplitudes and time constants. Starting with the model prediction on the evaluation batch, Table 1 shows the “Top 1” and “Top 3” accuracy values. Specifically, the Top 1 accuracy counts how often DLRN prediction and expectation have an exact match, while the Top 3 accuracy indicates when one of the three most probable predictions matches the ground truth.

**Table 1.** Accuracy of the predictions of kinetic model, time constants, and amplitudes from DLRN analysis of 100,000 2D data sets. A Top 3 accuracy value is also included for model prediction. Accuracy of time constants and amplitude calculated using the area metric *A*_M_.

| Spectra predictions | Accuracy |  |  |
| --- | --- | --- | --- |
| | Top 1 | Top 3 | $A_m$ |
| Model | 83.1% | 98.0% | – |
| Time const. ($A_m > 0.9$) | – | – | 80.8% |
| Time const. ($A_m > 0.8$) | – | – | 95.2% |
| Amplitudes ($A_m > 0.8$) | – | – | 81.4% |

Analyzing the evaluation batch using this accuracy measure revealed that in 83.1% of cases, the DLRN correctly predicted the expected model (with a prediction confidence of >95%), whereas in 14.9% of cases (which adds up to 98.0% for Top 3 in Table 1), one of the three most likely predictions matched the expected kinetic model. This shows that DLRN predicted the kinetic model of time-resolved spectral data sets with high confidence. For predicting time constants on the evaluation batch, DLRN accuracy was 80.8% using an area metric *A*_m_ > 0.9, which means that the forecasts have an average error of less than 10% of the expected values. If we reduced the area metrics to *A*_m_ > 0.8, 95.2% of DLRN predictions have an error of less than 20% of the original values. This indicates that DLRN almost always predicted time constants with an error of no more than 20% and were, in most cases, below 10%. High performance was also shown for the prediction of amplitude for the evaluation batch by DLRN. Using *A*_m_ > 0.8, the DLRN accuracy was 81.4%. However, DLRN extrapolates four spectra during the analysis for the amplitude prediction. The error limit (20%) is the sum of the errors of these four spectra, indicating that each spectrum is subject to an error of less than 5% compared to the expected outcome.

### DLRN analysis of time-resolved data having the first component as a dark state

Next, we wanted to test whether DLRN can also be used to analyze time-resolved data sets in which the initial state is a dark state, i.e., one that does not emit light but is involved in the dynamics. Whether all the components involved in the mechanism can be tracked depends on the experimental technique used to acquire the data. For example, to track enzymatic reactions, a dye is activated only if the enzyme interacts with a specific molecule, resulting in a rising dye signal that indicates the formation of new molecules induced by the enzyme population. Fig. 2a shows the expected model with a relative time constant for each decay pathway compared to the DLRN prediction. The DLRN analysis showed good agreement with the expected kinetic model. Similar performance was obtained for the amplitude prediction, which slightly differed for the expected spectra (Fig. 2b). Notably, the amplitude for the initial state A is equal to zero, indicating that DLRN can understand when the initial state is “off”, but that it can also extrapolate the correct kinetic model. In this case, DLRN was also tested on an evaluation batch of 100,000 2D data sets. The results were similar to those shown in Table 1. An approximate difference of only 1–2% was observed in the predictions of amplitudes and time constants, which were 79,5% and 94% for the time constant prediction with area metrics greater than 0.8 and 0.9, respectively, whereas DLRN accuracy for the amplitude analysis was 79.7% for *A*_M_ > 0.8. No significant changes were observed for the Top 1 and Top 3 model predictions, which showed similar accuracy values compared to the analysis performed on the batch dataset with initial state A “on”.

**Fig. 2.**
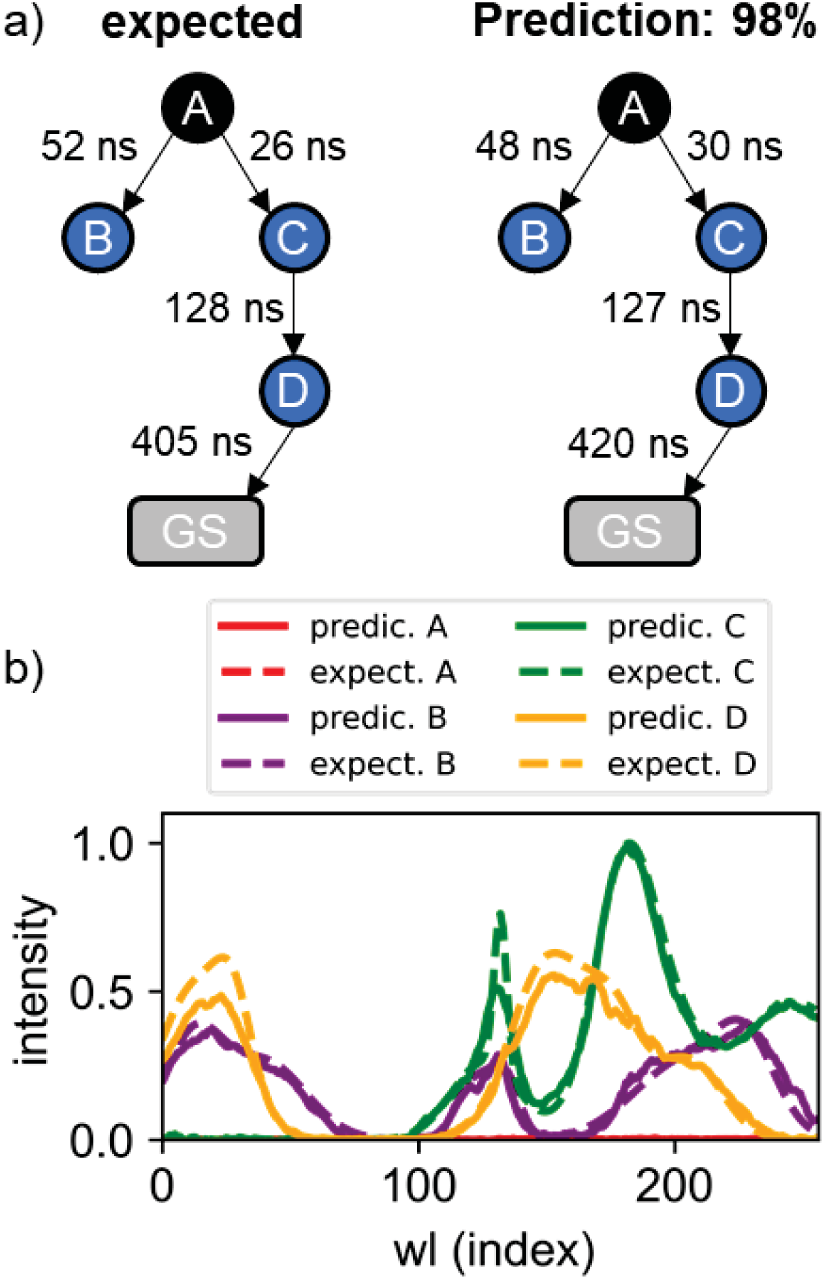
DLRN analysis of systems having an initial dark state. **a,** Expected and DLRN-predicted kinetic models with the corresponding time constant for each pathway. **b,** Comparison between expected (dashed lines) and predicted (continues lines) spectra associated with the kinetic model analyzed. As expected, DLRN predicts a null amplitude for the first dark state A.

### DLRN versus the classical method

The performance of the DLRN neural network was compared to the classical fitting analysis. Fig. 3a shows the expected model and relative spectra for the 2D data set used for the comparison. The time-resolved data set presents a complex mechanism, which is challenging to analyze. Starting with the model prediction, DLRN can identify the correct model with a probability confidence of 98.9% (Fig. 3b).

**Fig. 3.**
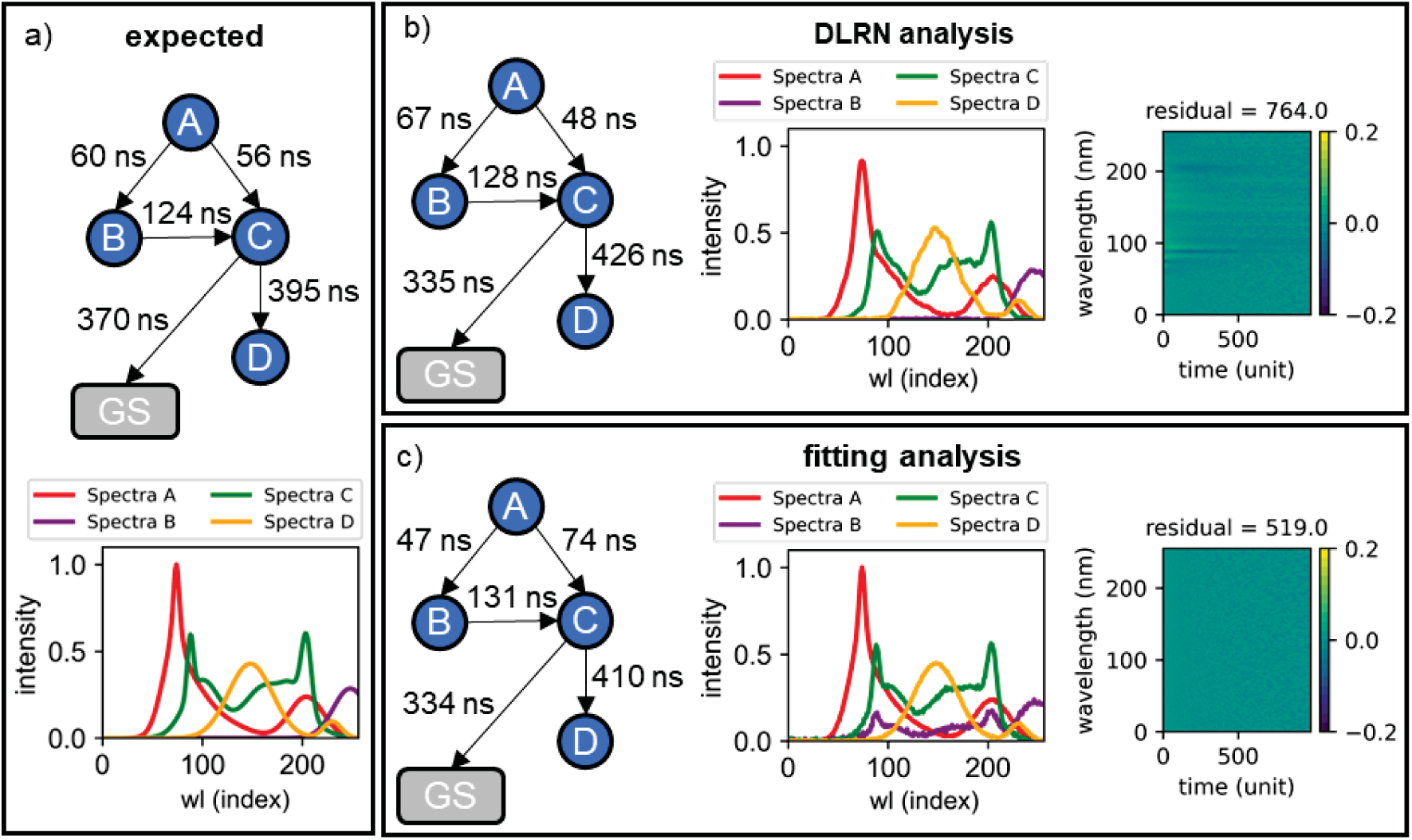
Comparison between DLRN and classical analysis used on the same complex molecular dynamics. **a,** Expected kinetic model with time constants for each decay pathway (top) and the respective spectra for each species (bottom). **b,** Kinetic model pathways and time constants (left), spectra for each species (middle), and residuals (right) obtained by DLRN analysis. **c,** Kinetic model pathways and time constants (left), spectra for each species (middle), and residuals (right) obtained by classical fitting analysis.

Moreover, the time constants predicted by the neural network match the expected one with good approximation, including the branching ratio (compare modeled time constants in Fig. 3a,b). Additionally, DLRN predicted accurately the amplitudes (Fig. 3b), showing that it can also extrapolate the correct spectra for complex dynamics. The residuals between the DLRN analysis and original data (Fig. 3b) are mostly uniformly distributed around zero, indicative of minor differences with the original signal. Fig. 3c shows the fitting analysis performed on the same datasets analyzed by DLRN. However, a classical analysis needs the kinetic model for the fit, whereas DLRN can extrapolate the model by itself. When the correct model was imposed for the classical analysis, the time constants and the amplitudes were free to change during the fitting for extrapolation. Starting with the time constants (Fig. 3c), the fit analysis shows a mismatch in the tau values, especially in the branching pathways of A (compare models in Fig. 3a,c).

Moreover, the amplitude of spectrum B shows several mismatches compared to the expected one (Fig. 3b, purple line), whereas the others are comparable. This mismatch can be attributed to compensating the time constants during the fitting, indicating that a local minimum was reached during the analysis. Residuals of the fit analysis (Fig. 3b) are better than those of DLRN (Fig. 3c), but the differences were not significant.

During the classical analysis, we varied the initial parameters to further improve the fit of the analysis. Here, we observed three solutions having the same residual score, similar to that shown in Fig. 3c, indicating that more than one minimum was obtained during the analysis. Depending on the minima, time constants and amplitudes can be more compatible with those expected (Fig. SI 7). In the most straightforward cases, DLRN analysis and classical fit analysis converge, on average, to similar or even the same results (Fig. SI 6). The results presented in this section show that DLRN can perform at least as well as classical fitting and, in some cases, even better. It can obtain the correct minimum solution over multiple local solutions, and automatically extrapolate all parameters for the analysis in a few passes. This allows fast and accurate computation of time-resolved data.

### Multi-timescale data analysis

Next, DLRN was used to analyze data sets having two orders of magnitude to test if the output of the neural network can be used to obtain kinetic information for systems that evolve on more than one timescale. We generated three datasets on the millisecond (first order) and microsecond (second order) timescales with random kinetic model, time constants, and spectra. The dynamics of each dataset are summarized in Fig. 4a–c, where active states and their corresponding decay pathways and time constants are shown. Each data set was divided into two subsets, corresponding to each timescale, and interpolated to match the timescale used for training DLRN. After that, each subset was analyzed using DLRN, and the results combined to obtain the total kinetic model (Fig. 4d–f), as is commonly done in a classical analysis.

**Fig. 4.**
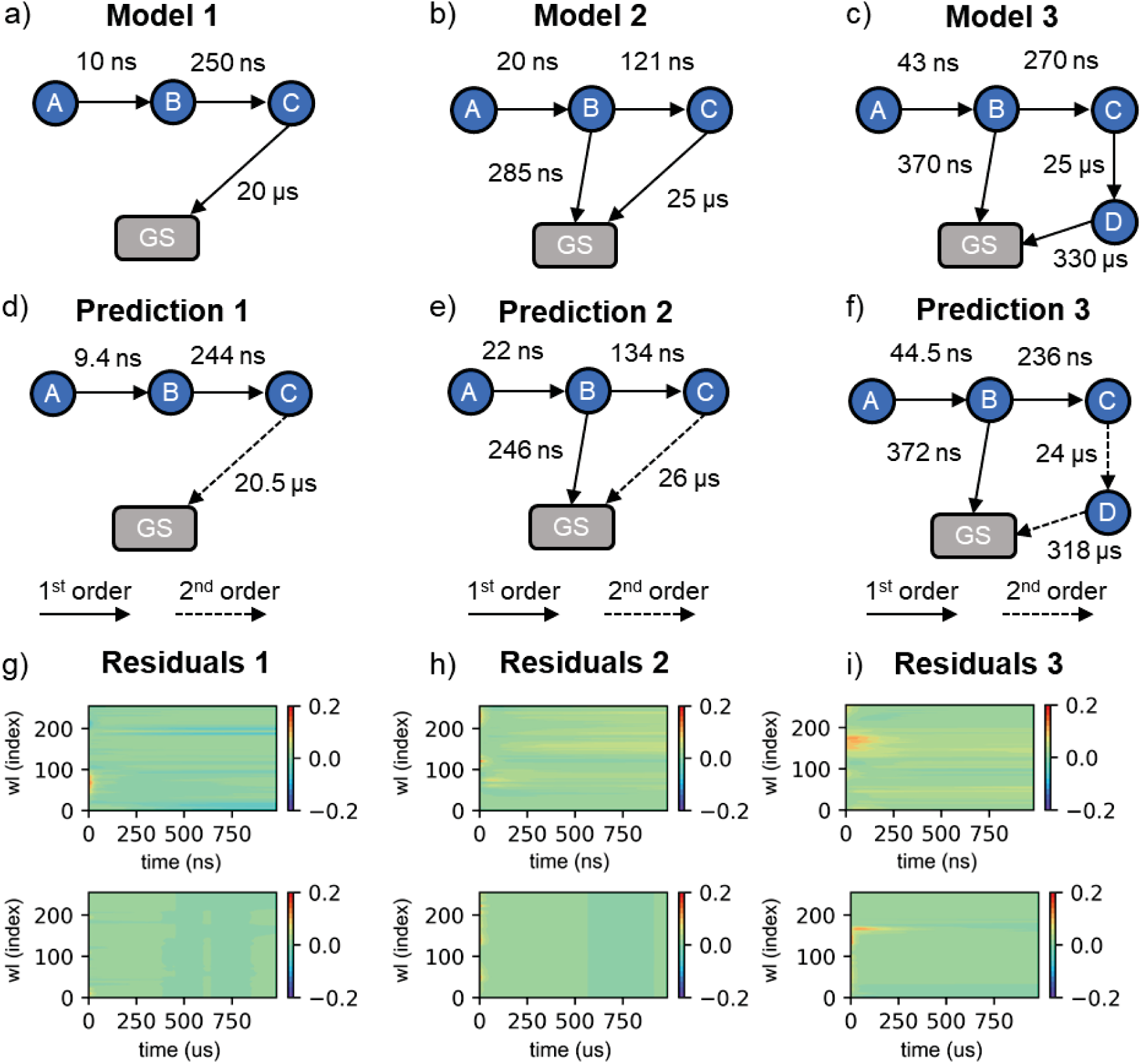
DLRN reconstruction analysis on a multi-timescale problem. **a–c,** Kinetic model used for generating the first (a), second (b), and third (c) data sets for the multi-scale analysis. **d–f,** Kinetic model prediction obtained using DLRN for the first (d), second (e), and third (f) data sets. **g–i,** Residuals obtained by fitting the two orders of magnitude subsets using the concentration matrix **C_DLRN_**(t) and spectrum matrix **S_DLRN_**(λ) obtained by DLRN for each multi-scale dataset.

Additionally, the residuals of the fit using DLRN are shown for the two subsets for each system analyzed (Fig. 4g–i). To obtain the residuals, the concentration matrix **C_DLRN_**(t) corresponding to the DLRN model prediction was multiplied by the spectrum matrix **S_DLRN_**(λ), which DLRN also obtains, and the result was subtracted from the subset for each timescale. Starting with the neural network prediction on the first model (Fig. 4d), DLRN extrapolates a linear kinetic model having three active states (A, B, and C), two of which decay on the nanosecond timescale (first-order analysis) with time constants *T_A_* = 9.4 ns and *T_B_* = 244 ns, while one decays on the microsecond timescale with *T_e_* = 20.5 μs (second-order analysis). After comparing these results with the expected kinetics, we conclude that DLRN correctly predicted the dynamics in this case. Moreover, the excellent quality of the residuals (Fig. 4g) shows that the spectrum matrix **S_DLRN_**(λ) was also correctly extrapolated by DLRN. For the kinetic models of the other two systems (Fig. 4b,c), the data show more complex dynamics with branching pathways and more electronic states to extrapolate. Despite this, DLRN correctly predicted the expected models with good approximation of the time constants (Fig. 4e,f). Furthermore, the residuals show good values (Fig. 4h,i), indicating that the spectrum matrix **S_DLRN_**(λ) obtained in these cases is plausible.

### Analysis of synthetic time-resolved agarose gel electrophoresis using DLRN

The DLRN was then trained to analyze synthetic images of a time-resolved agarose gel. These gels are used to track molecular changes at different time delays. We were interested in using DLRN to obtain and investigate the molecular dynamics that control such systems. The synthetic agarose gel images were generated similarly to the spectral images, with the difference that the gels had a maximum number of 17 time points (the possible cells of the gel), which means that each time point covers more x pixels. The base pair (Bp) spectrum of each state was generated by summing a random number of one to eight Gaussian functions with random peak position (between 0 and 256 every 2 indexes), width (between 2 and 10), and intensity (between 0.2 and 1). Each Bp spectrum has at least one amplitude between 0.4 and 1. Fig. 5a shows an example of a time-resolved agarose gel analysis. The results of DLRN analysis of 100,000 2D data of time-resolved agarose gel are shown in Table 2. DLRN model prediction had an accuracy of 82.4 % for the Top 1, which means that in these cases, DLRN correctly predicted the expected model with high-probability confidence. In 16.2% of cases, i.e., a total accuracy of 98.2% for the Top 3, one of the three most likely predictions obtained from DLRN matches the expected kinetic model. DLRN showed good performance in predicting time constants. In 72.8% of cases (*A*_m_ ≥ 0.9; Table 2), we obtained an error of less than 10%, while in 92.1% of the predictions (*A*_m_ > 0.8; Table 2), the error was less than 20%.

**Fig. 5.**
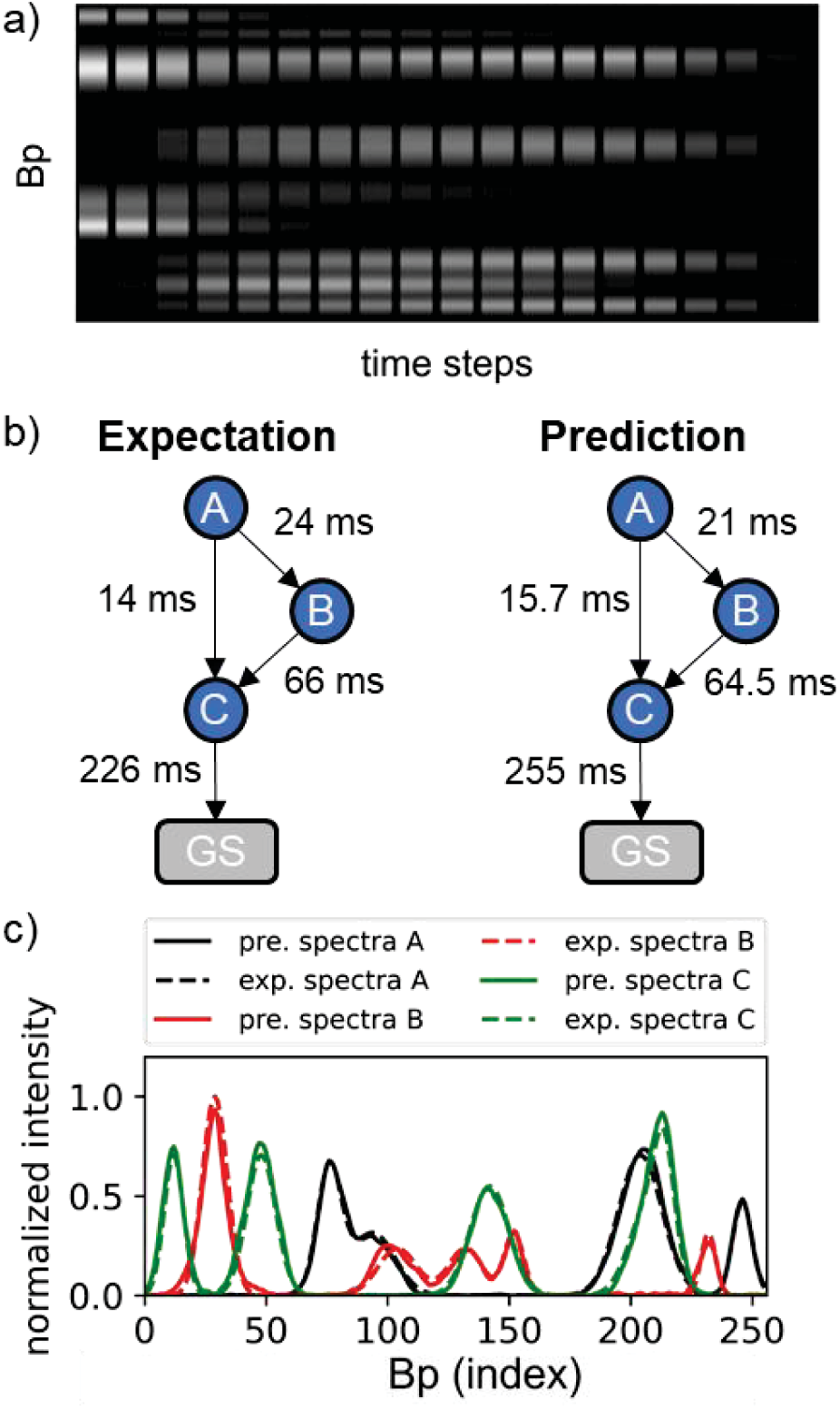
DLRN analysis of 2D time-resolved agarose gel electrophoresis. **a,** Example of a synthetic agarose gel used for the kinetic model analysis with DLRN. **b,** Expected model and DLRN analysis performed on the image shown in panel a. DLRN can correctly predict both the kinetic model and the time constant associated with each pathway. **c,** Comparison between DLRN prediction (continuous lines) and expected (dashed lines) amplitude from the image in panel a.

**Table 2.**
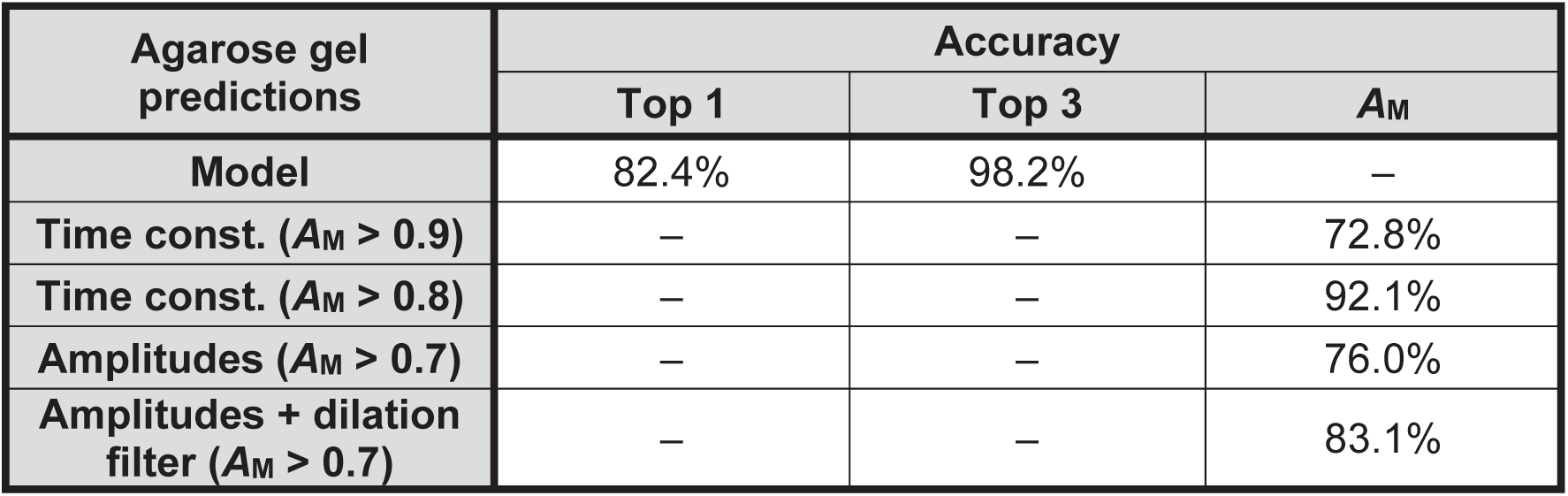
Accuracy of the predictions of kinetic model, time constants, and amplitudes from DLRN analysis of 100,000 2D data sets (agarose gels). A Top 3 accuracy value is also included for model prediction. Accuracy of time constants and amplitude calculated using the area metric *A*_M_.

The amplitudes of agarose gel spectra were also analyzed using DLRN. Here, predictions of individual amplitudes differed on average by no more than 7% from the real spectra in 76% of the batch dataset. We observed that this percentage could be increased to 83.1% by adding a dilation filter (= 3) in all the separable convolution 1D layers in the second residual block (see amplitude block, Fig. SI 2). These high-performance results show that DLRN can also be used to analyze time-resolved agarose gels for tracking dynamics during molecular assembly. Further DLRN analysis and results of time-resolved agarose gels are reported in the Supplementary Information (section 5).

### DLRN analysis on DNA strand displacement networks

We used DLRN to analyze and disentangle DNA strand displacement (DSD) CRNs. DSD CRNs provide an efficient toolbox with many design options (network architecture, reaction rates) to create complex nonlinear behavior in life-like materials systems based on the strict rules of DNA base-pairing. The kinetics of simple DNA strand interactions can be predicted very well numerically (e.g., using the Visual DSD program^46^). However, these numerical predictions are no longer realistic if enzymatic reaction steps, pH-dependent conformational changes, or the presence of dye and quencher molecules on the DNA strands affect the kinetics of DSD. In these cases, DLRN analysis is necessary to understand the dynamics of a complex DSD CRN. Here, DSD dynamics involve four different DNA strands (Fig. 6a), denoted input (Inp), substrate 1 (Su1), substrate 2 (Su2), and reporter (Re). The mechanism of DSD is a branching system in which Inp (state A) can interact with Su1 and Su2, liberating two different fluorescent strands that correspond to states B and C. Then, these two strands can react with Re, and both liberate the same last fluorescent DNA strand (state D). We designed versions of the CRN with different sequences (sequence fragment 1 in blue of Su1, Fig. 6a) to tune the reaction rate of branch C. The corresponding DNA sequence is given in the Supplementary Information (section 8). Limited to pseudo-first-order reactions, DLRN was initially used to analyze the simple cases in which one of the branching pathways was turned off by using only one substrate at a time. Data generation is described in detail in the Methods section. Fig. 6 shows the comparison between the DLRN analysis and the expected model, spectra, and kinetic traces obtained by Visual DSD simulations only if Su1 = 20 nM (Fig. 6b–d) or Su2 = 20 nM (Fig. 6e–g); Inp and Re concentrations were constant at 10 and 30 nM, respectively. In both cases, DLRN was able to correctly identify the CRN (Fig. 6b,e), extrapolate the correct spectral amplitudes involved (Fig. 6c,f), and reconstruct the kinetic traces (Fig. 6d,g). Although expected (Visual DSD) and predicted (DLRN) kinetic traces overlap with each other, the values of *T_e→D_* and *T_B→D_* differed significantly from the expected values (in particular, the *T_B→D_* value differed by about 35%), while the values of *T_A→B_* and *T_A→e_* match the values with an error of less than 10%. Due to this discrepancy, we created the kinetic traces for both the dynamics imposing for *T_e→D_*, and *T_B→D_*. The calculated values were obtained by multiplying the expected rate constant *k_e→D_* and *k_B→D_* from visual DSD by the Re concentration. This was followed by comparing the forced and expected (Visual DSD) kinetic traces (Fig. SI 13). The comparison showed that the forced and predicted kinetic traces diverged significantly, indicating that the DLRN analysis was correct. Therefore, the discrepancy between the expected and predicted tau values is not associated with the neural network’s performance but with possible non-first-order kinetic mechanisms.

**Fig. 6.**
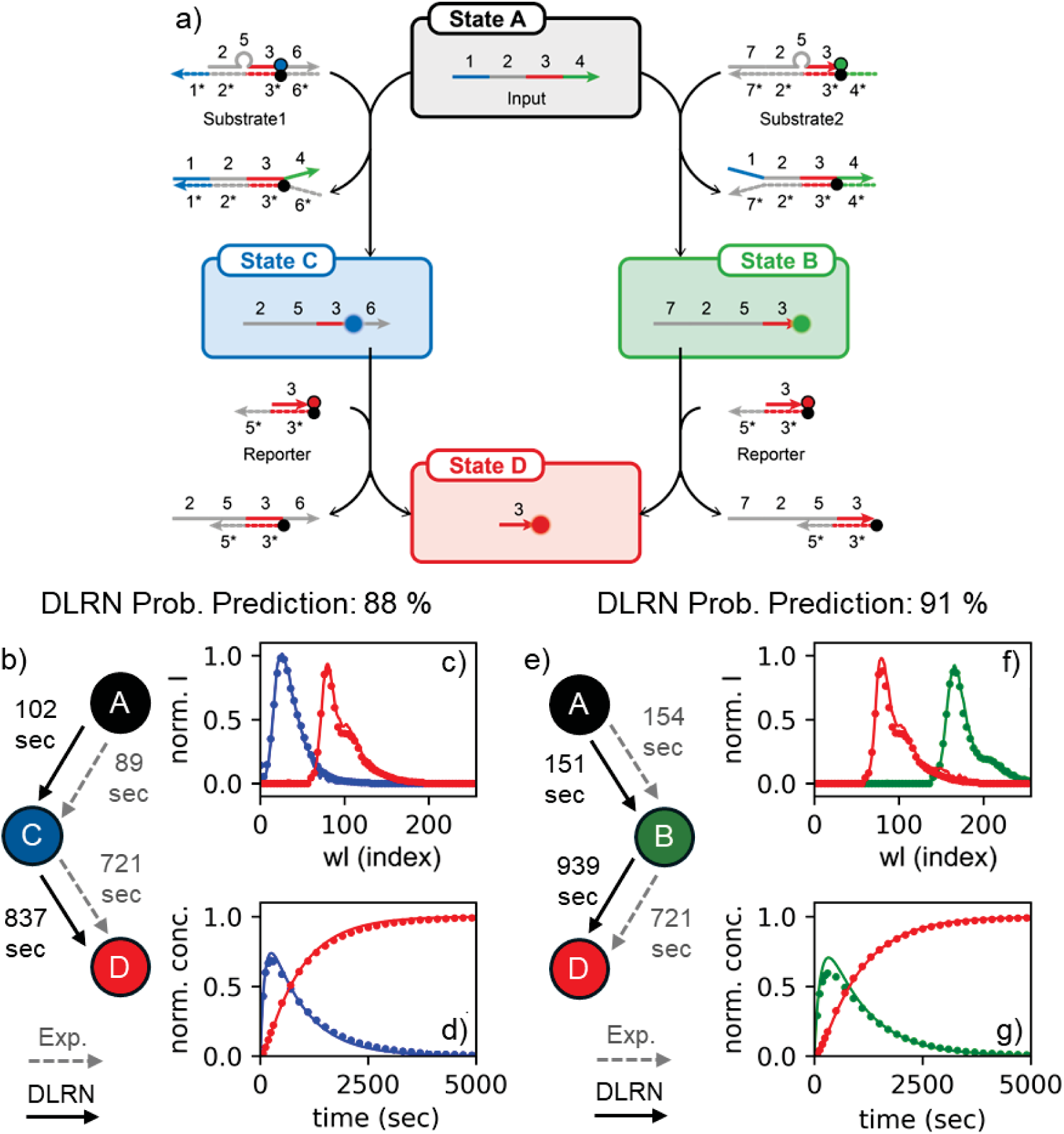
DSD kinetic reaction analysis for Su1- and Su2-only dynamics. **a,** General DSD kinetic reaction mechanism. Region 1 (blue continue line in the input vector of state A) is the toehold. **b** and **e,** Comparison between DLRN (continuous black arrows) and expected (dashed gray arrows). Kinetic models for Su1 only (b) and Su2 only (e) with time constants shown for each decay pathway. The initial state A is a dark state. **c** and **f,** Expected (filled circles) and DLRN-predicted (continuous line) spectra for Su1 only (c) and Su2 only (f) dynamics. **d** and **g,** Expected (filled circles) and DLRN-predicted (continuous lines) kinetic traces for Su1 only (d) and Su2 only (g) dynamics. Colors correspond to the kinetic model.

This discrepancy indicates that the B→D and C→D pathways have more complex dynamics and that multiplying the expected rate constants by the Re concentration is not sufficient to calculate the correct time constants. To extrapolate the correct rate constant, DLRN was used to analyze four data sets with different Re concentrations. After DLRN analysis, the inverse of *T_e→D_* obtained from DLRN was plotted versus Re concentration and then the data fitted using a linear regression (Fig. SI 14). Using this method, second-order reaction pathways can be tracked and the expected rate constant *k*_0_ (intercept from the linear fitting) can be calculated. The intercept value obtained from the linear regression is 4.83 × 10^−5^, which differs from the expected value, *k*_CD_ = 4.62 × 10^−5^, by approximately 4.5%.

The DLRN was also used to analyze the case in which both Su1 and Su2 are present and can interact with the input strand, thus generating a branching mechanism for the formation of the final state D. Fig. 7 shows the results obtained from DLRN analysis using Inp = 10 nM, Su1 = 12 nM, Su2 = 8 nM, and Re = 30 nM. The neural network was able to establish the correct network architecture with a probability of 75% (Fig. 7a), and correctly reproduced the kinetics of the dynamics (Fig. 7c), indicating that the predicted tau values can represent each decay pathway. However, the amplitude was not well disentangled from the analysis in which the spectra of the states C and B mixed with each other. Despite differences in expected amplitude, residuals from DLRN analysis were good (Fig. SI 15a), indicating a correct analysis of the dynamics. This behavior can be attributed to the identical rising-decaying kinetics of the states C and B (Fig. SI 15b), which makes them extremely difficult to disentangle. We also tried to analyze the data by classical fitting using a branching mechanism. For the fit, a branch coefficient between 0.01 and 1 for state A, and positive amplitudes were imposed. Surprisingly, the DSD CRN was better analyzed by DLRN than by classical fitting (Fig. 7 and Fig. SI 16). Kinetic traces and time constants were better resolved by DLRN, whereas classically fitted traces deviated significantly from those expected. Conversely, classical fitting extrapolated cleaner and separate amplitudes. However, the spectrum of B differed greatly from the expected position, which was not the case with DLRN.

**Fig. 7.**
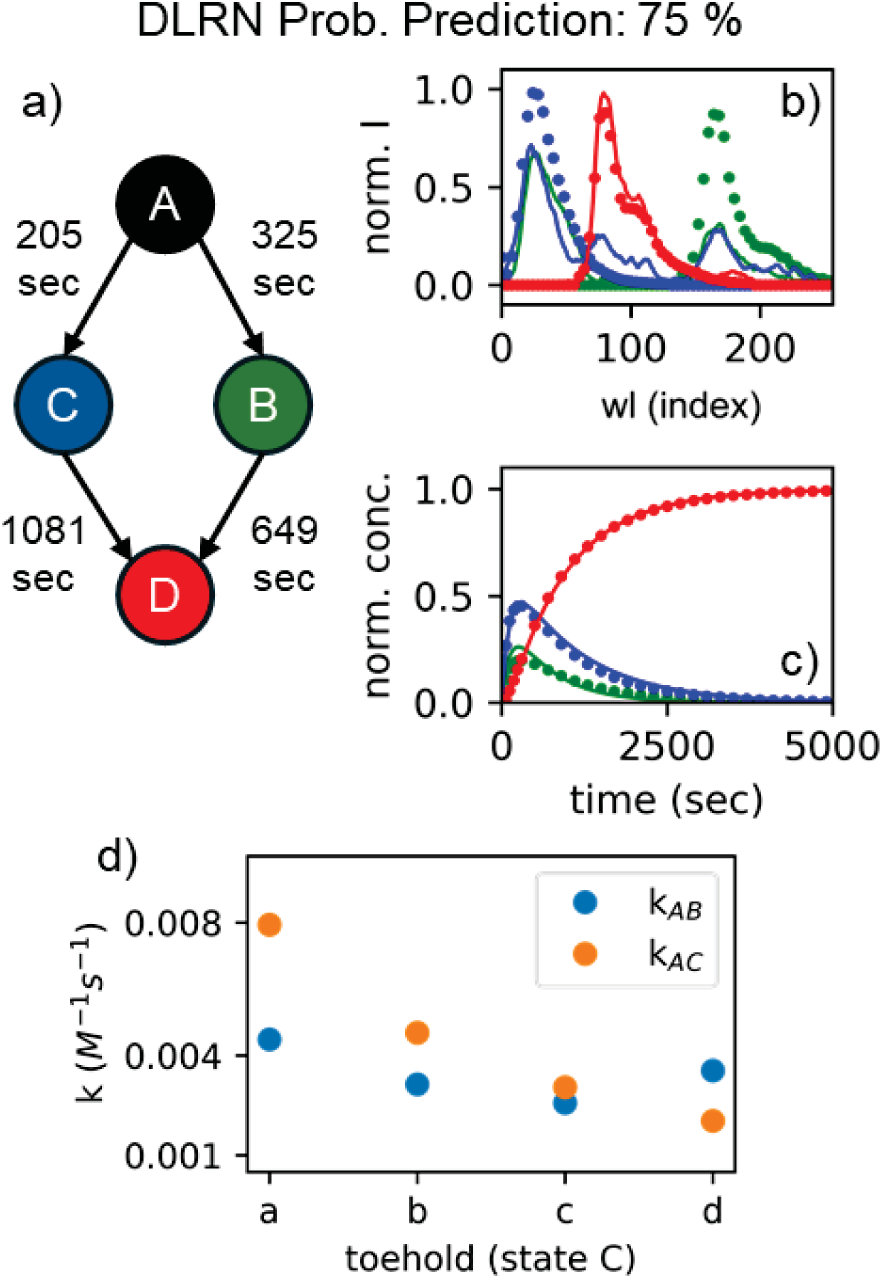
Full DSD kinetic reaction analysis. **a,** DLRN model prediction. **b,** Expected (filled circles) and DLRN-predicted (continue line) spectra for the full DSD dynamics. Colors correspond to the kinetic model in panel a. **c,** Expected (filled circles) and DLRN-predicted (continuous lines) kinetic traces for DSD dynamics. Colors correspond to the kinetic model in panel a. **d,** Rate constants *k*_AB_ (blue) and *k*_AC_ (orange) obtained from DLRN analysis of four variant Input strands by changing the toehold of the DNA strand.

We analyzed the DSD dynamics, which had a series of different Inp strands with varying toehold (varied domain 1 of the Inp DNA strand, blue sequence Table SI 2). Because the corresponding toehold mediates the reaction rate of each DSD step, this modification affects the formation of state C. The longer/stronger the toehold binding, the faster the migration of the whole DNA-branch from Inp to Su1. We designed four different DNA toeholds that decelerate the kinetic pathway A→C, and extrapolated the rate constants *k*_AC_ and *k*_AB_ using DLRN. The concentrations of Inp, Su1, Su2, and Re were kept constant at 10 nM, 10 nM, 10 nM, and 30 nM, respectively, and the results are shown in Fig. 7e. DLRN was able to track the variation of *k*_AC_ for the toeholds used, predict the correct branching mechanism, and follow the expected kinetic traces (see also Fig. SI 17). The kinetic pathway from state A depends on *k*_AB_ and *k*_AC_ values, and it was possible, especially from the kinetics traces, to observe preferences. With toehold a, population of state C is preferred, whereas for toehold d, decay into B is preferred.

### G-User-DLRN: a graphic user interface for DLRN analysis

To increase the method’s usability by obtaining a DLRN analysis in a few concise steps, a Python-based graphical user interface (GUI) was developed (G-User-DLRN). After running the file “DLRN_GUI”, a graphical interface starts, and several options are available for the user to select to begin the analysis. The first option allows the user to use the pre-trained DLRN model for spectral or agarose gel analysis. The button “load custom scale” loads the .txt file containing a vector of time points used for the measurements. This allows the rescaling of data to match the timescale used to train the DLRN. The loaded data is shown in an additional window after selection via the “load data” button. The GUI also allows the Top 1 and Top 3 analysis outputs to be chosen. After these selections are made, clicking the “data analysis” button generates a number of windows depending on whether Top 1 or Top 3 was chosen; each window shows the complete analysis for the most probable (Top 1) or the three most probable solutions (Top 3). Each window shows a kinetic model representation with the DLRN probability, an indication of how confident the DLRN is regarding a particular outcome.

Additionally, tau, amplitude, and population profiles obtained by DLRN analysis are shown in the window for each prediction. Moreover, residuals normalized by the maximum value of the dataset are presented in the window to understand how representative the predictions are of the data. The analysis also provides a table showing the time constant value predicted by DLRN and the model prediction in one-hot encoding representation. At the end, G-User-DLRN has a button called “test DLRN”, which performs the analysis on test data, but with the difference that expected values for model, tau and amplitudes are shown in the result’s windows. This option can be used to check the performance of DLRN on ground truth data sets. Additional information, including a primary pipeline and images of G-User-DLRN can be found in the Supplementary Information (section 7).

## Discussion

We have introduced DLRN, a deep learning framework for the GTA of 2D time-resolved data sets. The standard approach of fitting kinetics using GTA is almost compulsory for the analysis of dynamic systems because it can quantitatively extrapolate kinetic mechanisms from time-resolved data^5,15–18^. However, the path toward achieving results can be long and complex, requiring a lot of time and an interwoven approach of mathematical modeling, assumptions, and experimental data. Furthermore, even with a good approach that guides one through the analysis, a family of models might show the same residual performance and reasonable results, making it difficult to distinguish from the analysis alone and requiring further work to identify the “best” solution. Indeed, this was the case during the classical fitting test (Fig. 3 and Fig. SI 7), where the global and target fitting gave suitable residual and reasonable model parameters, but several global minima were identified. The GTA solutions depended on the initial fit parameters, making it difficult to identify the correct solution for our system.

Moreover, the kinetic model was already known and used for the analysis, which drastically reduces the workload for a classical analysis by excluding all the other possible models that can fit the data. Conversely, DLRN alone could extrapolate the correct kinetic model with a confidence of 98.9% (Fig. 3b). Moreover, it could correctly conclude both the time constants and the amplitude shape (Fig. 3b). Classical fitting was only superior in terms of residual scores, which were higher with DLRN, although the two were somewhat comparable in this aspect. This arises because the DLRN neural network yields a fixed solution with a margin of error. Indeed, the analysis performed on a test batch showed excellent performance (Table 1) but with some flexibility. One of the critical roles of flexibility lies in the Top 1 and Top 3 solutions. The performance of DLRN shows that the neural network can become confused and not clearly distinguish one solution from another. However, it gives a maximum of three possible models (each has its amplitude and time constants), and the probability that the real solution is one of these is greater than 98%. At this point, the probability confidence (y value in Fig. 1b) can be combined with the residuals between the DLRN results and data to obtain the most probable solution that best reflects the system. The speed of DLRN is remarkable—it is capable of extrapolating a complete analysis in seconds.

An exciting aspect of DLRN is that it can be used to analyze more than one type of time-resolved data. Despite spectral evolution, DLRN was used to analyze time-resolved agarose gel electropherograms (Fig. 5). Compared to the spectral analysis, DLRN performance with gels was lower (compare Tables 1 and 2); however, the results were still good (Figs. SI 10–12). The most noticeable differences were observed for the predictions of time constants and amplitudes, while model prediction differed by less than 2% in both Top 1 and Top 3 values compared to the spectral analysis. Differences in tau and amplitude values can be expected due to the nature of the agarose gel, which is made up of cells, thus reducing the number of actual points carrying dynamics information.

Another crucial point was to evaluate DLRN for multi-timescale analysis (Fig. 4). The neural network was used to analyze a signal that evolves into two timescales, from the nanosecond to the microsecond. The signal was decomposed into two data sets, each with only one timescale, and subsequently analyzed by DLRN. DLRN could also extrapolate the correct kinetic model and its parameters. This shows that DLRN can be used to analyze data sets on more than one timescale. Moreover, DLRN architecture is thought to extrapolate four species for each timescale. This means that if more than four species are involved in the dynamics but split over several timescales, DLRN can theoretically extrapolate all of them by analyzing each time window individually.

DLRN was used to analyze DSD CRNs in different conditions to emulate its utility for complex systems under certain conditions. The expense of DNA limits the number and type of experiments from which one can collect data for classical fitting. In addition, classical methods cannot predict certain effects (e.g., enzymatic steps or pH-dependence). Although DSD reactions are usually second-order, for both linear (Fig. 6) and branching (Fig. 7) mechanisms, DLRN was able to correctly reproduce the DSD kinetics (pseudo-first-order-reaction limited), predicting kinetic model, time constants, and amplitudes of the systems. DLRN was also compared to the classical fit for the branching mechanism (the most complex one), and surprisingly, the neural network could better reproduce the system’s dynamics (Fig. 7 and Fig. SI 16). Moreover, using DLRN, we were able to extrapolate the correct rate constants *k*_CD_ within an error of 4.5% (Fig. SI 14) and the kinetics for four different DNA toeholds for the full dynamics with Su1 and Su2 both present (Fig. 7e). Together, these results demonstrate the efficiency of DLRN in kinetic analysis and its versality for systems of varying complexity.

## Materials and Methods

### Neural network architecture

The core methodological component proposed here is a combination of an architecture based on Inception-Resnet^45^, which is composed of four blocks (stem, ConV. A, ConV. B, and ConV. C) and three personalized output blocks (model, time, and spectrum). This approach created a deep learning reaction network (DLRN) for analyzing 2D time-resolved data sets, as schematically visualized in Fig. SI 1. A more detailed representation architecture is shown in Fig. SI 1. As a general overview, each Inception block comprises several convolutional and pooling layers in parallel, where kernel size, filters, and resize depends on the specific block. Each block is repeated in the network architecture several times: once for stem, five for block A, 10 for block B, and five for block C.

Additionally, two types of reduction modules are introduced after the total repetitions for both A and B blocks. In the case of the output blocks, the architecture changes depending on the desired output (Fig. SI 2). The model block determines the kinetic model that best represents the data over 102 possible kinetic models. It comprises an averaging pool layer, a dropout layer, and a dense layer with a Softmax activation function (Fig. SI 2a). The categorical cross-entropy loss function is used to optimize the model prediction. The time block was designed to extrapolate the kinetic time constants associated with the kinetic pathways of the obtained model. This block has two inputs: the output from the last block C, and the concentration matrix linked to the specific kinetic model in the analysis. The two inputs are processed in parallel using several dense layers and residual convolutional blocks to obtain the number and the value of the time constants involved in the dynamics using a regression problem (Fig. SI 2b). The mean squared error was used as a loss function for the time block. The amplitude block extrapolates the number and the shape of the species involved in the dynamics, which are linked to the amplitudes of each decay component that contributes to the signal. Similar to the time block, the amplitude block also has two inputs: the output from the last block C and the concentration matrix (see also Fig. SI 2C). The two inputs are initially analyzed in parallel and then merged and sent through several separable convolutional and convolutional layers. As with the time block, the linear activation function was used for the last layer, and the mean squared error was applied as a loss function. The neural network was trained using 2.3 million synthetic images each with a size of 256 × 256 × 1. The Adam optimizer was used for the training with a starting learning rate of 0.0001.

### Generation of synthetic data for training and evaluation

To obtain an optimal and comprehensive data set, and due to the limited experimental data available, we used synthetic data to train and evaluate the neural network. For this purpose, a kinetic model with five electronic states (four excited states and one ground state) was randomly created to produce the respective synthetic data sets. However, some constraints were applied to the generation of the model, such that only physically feasible models could be generated (a total of 102). The assumptions used to reduce the complexity of the prediction task were: 1) there is only one initial populated state; 2) a forward reaction without cycling; 3) each active state can only populate a maximum of two other states. Restrictions 1 and 2 are physically motivated since perturbed states are not in equilibrium, and only one molecule is usually initially triggered. Assumption 3 is made for practical reasons to reduce the total number of possible models. Moreover, it is decidedly challenging to distinguish branching between three or more states in kinetic reactions, so the third assumption is a reasonable restriction.

The data sets for training and evaluation were generated by simulating time-resolved data **X**(t, λ) for each physical kinetic model. Two components are necessary to create this data: the active spectra matrix **S**(λ) of the electronic state and the concentration trace matrix **C**(t) of the selected kinetic model. The product **C**(t)\***S**(λ) generates the 2D signal **X**(t, λ), which is the sum of all the spectra at each time delay. Spectra of each state **S_i_**(λ) were generated by summing a random number of one to eight Gaussian functions with random peak position (between 0 and 256 every 2 indexes), width (between 3 and 26), and intensity (between 0.2 and 1). Notably, at least one peak of each spectrum has a minimum intensity of 0.4. This allows generation of sequential spectra with complex structures from the initial to the last populated state. The concentration matrix **C**(t) was created by converting the kinetic matrix associated with the randomly selected kinetic model into differential equations using a fifth-order Runge–Kutta method. The time constants for active pathways were randomly generated in sequence (from *T_1_* up to *T_n_ = 7*) from a predetermined nonlinear scale of approximately 210 values (from 1 to 605). The time points were interpolated by combining a linear growth scale with an exponential growth scale to obtain a total of 256 time points.

### Generation of DSD data for testing the DLRN

The DNA sequences of the domains were designed intuitively, and their crosstalk was avoided by checking their interactions using NUPACK^47^. The stem sequences are gray, the toeholds are colored (Table SI 1). Table SI 1 shows the exact DNA sequences with the fluorophores and quenchers. The rate coefficients of binding and unbinding of complementary DNA sequences were derived as shown in the Supplementary Information (section 8), and the values are summarized in Table SI 2.

### Area metric for regression accuracy

To determine the accuracy and power of the regression analysis, we introduced the measure of area metric *A*_M_. This measure is calculated from the ratio between the residuals and the areas of the expected values (Eq. 1). Because Δx of the data is constant, the ratio can simply be expressed as the sum of the residuals (**Res**) divided by the sum of the expected values (**ExpVal**).

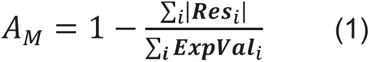

*A*_M_ is a value between 0 and 1, where 1 indicates that the two areas overlap entirely. By setting a lower bound for good area overlap, it is possible to determine how many predictions of the entire batch can be considered a positive result. This reduces the problem to a binary one, where accuracy can be calculated by dividing the number of positive predictions *N*_p_ by the total number of samples *N*_tot_ (Eq. 2):

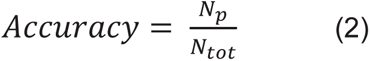

Throughout this study, we set the lower limit on an excellent area overlap at 0.8 and 0.9 for the predictions of time constants and 0.8 for amplitude predictions (the choice of these limits is explained in the Supplementary Information, section 4).

### Code availability

The code is available at https://github.com/mem3nto0/DLRN_App.git

