## Supplemental Information for "Deep Learning Reaction Network: a machine learning framework for modeling time resolved data"

#### 1. Deep learning reaction network (DLRN) architecture

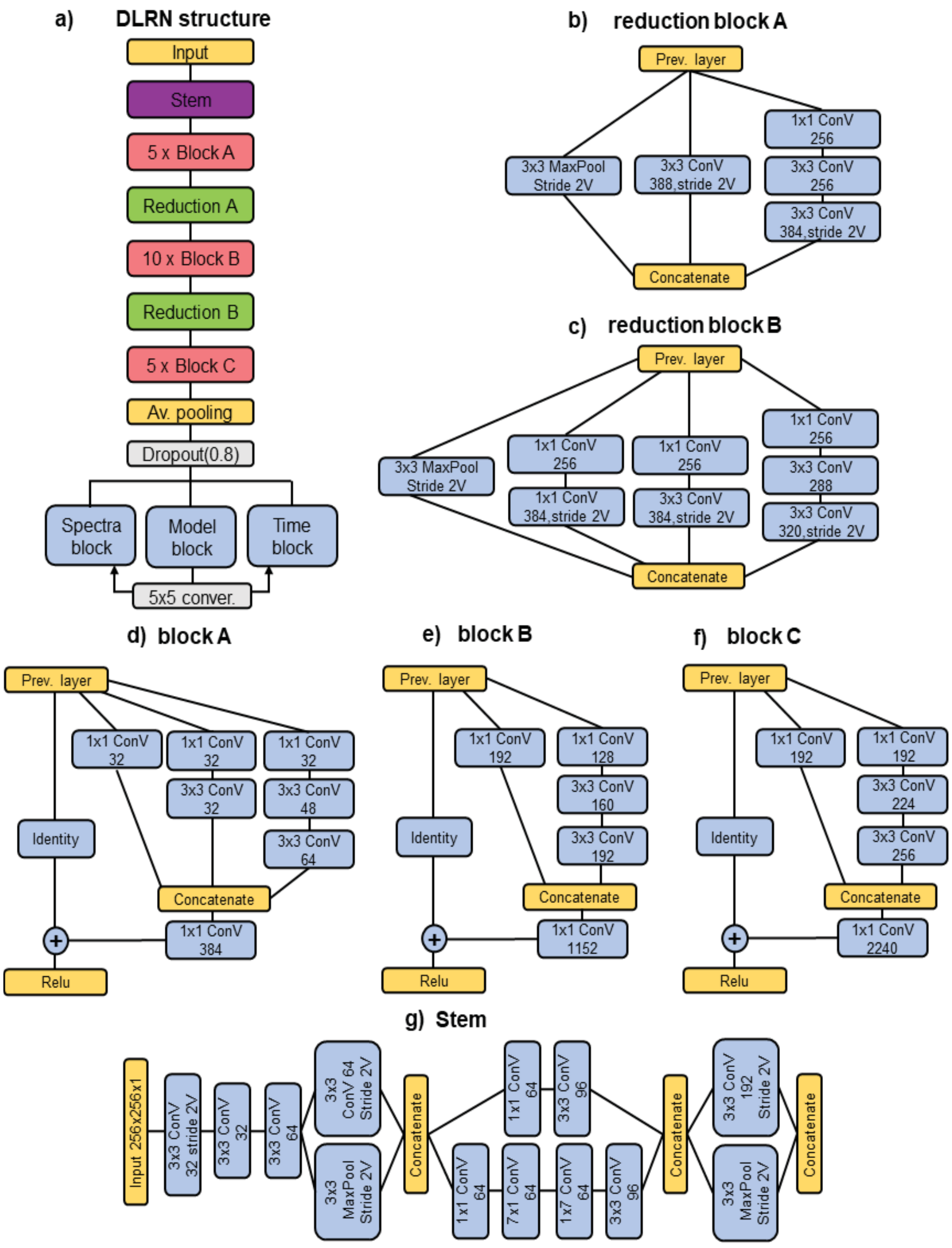


**Fig. SI 1. a,** Full architecture of the DLRN neural network. **b,** Reduction block after A blocks. **c,** Reduction block after B blocks. **d,** Residual block A. **e,** Residual block B. **f,** Residual block C. **g,** Stem block.

Fig. SI 1 shows the entire architecture of the DLRN. Input data—images with dimensions 256×256×1; time and wavelength in the x and y axes, respectively—are sent through the neural network to extrapolate the model, time constants, and spectra from the 2D data. Model, time, and spectra blocks are shown in Fig. SI 2.

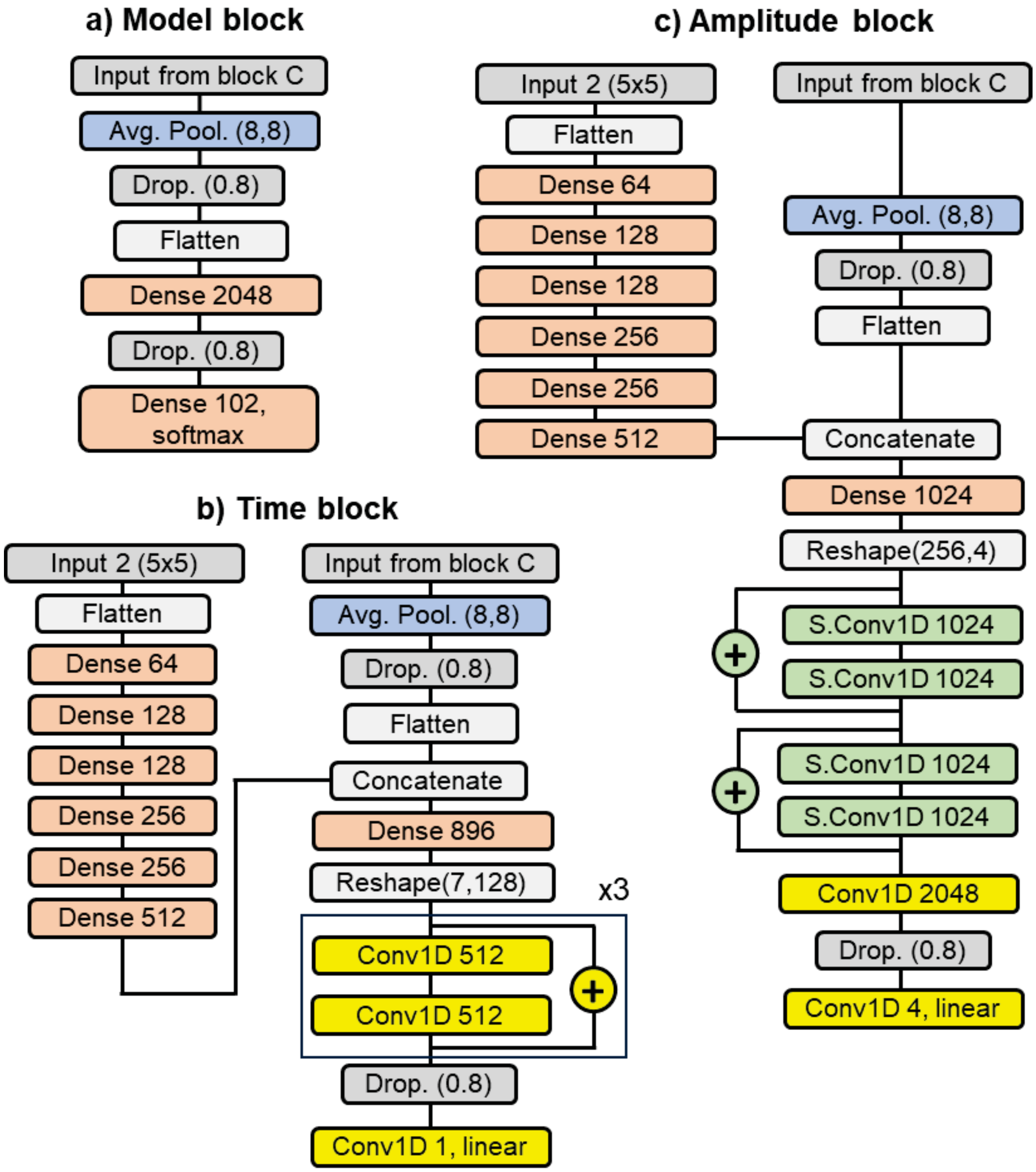


**Fig. SI 2. a,** Model block. Input from the last block C is sent through averaging and a dense layer, which uses Softmax activation. **b,** Time block. The model block provides one of the inputs after conversion in a concentration matrix reference. In parallel, block C's output is a second input. The two inputs are processed and then concatenated and sent through three residual convolutional blocks. Linear activation is used for the last convolutional block after dropout. **b,** Amplitude block. As for the time block, the two inputs are converted outputs from the model block and block C. To obtain the amplitudes, the two inputs are elaborated using dense, separable convolution 1D

(S.Conv1D), and convolution (Conv1D) layers. A linear activation function is used in the last layer.

### 2. Evaluating DLRN analysis: model, time constants, and amplitude

Here, we compare the DLRN prediction to the expected results for some test data sets. Fig. SI 3 shows a comparison of DLRN-predicted models (red lines) and expected models (black dashed lines) for 10 random data sets. Specifically, the x position of each plot indicates the model index, which is linked to a specific concentration trace matrix **C**(t) of a kinetic model. The y axis represents the confidence probability. DLRN-predicted and expected models overlap excellently, with prediction confidence (y value) greater than 98% in almost all examples. A few cases showed multiple outputs from the DRLN analysis. However, at least one of the top 3 DLRN solutions matched the expected one, indicating that one possible solution is the exact model.

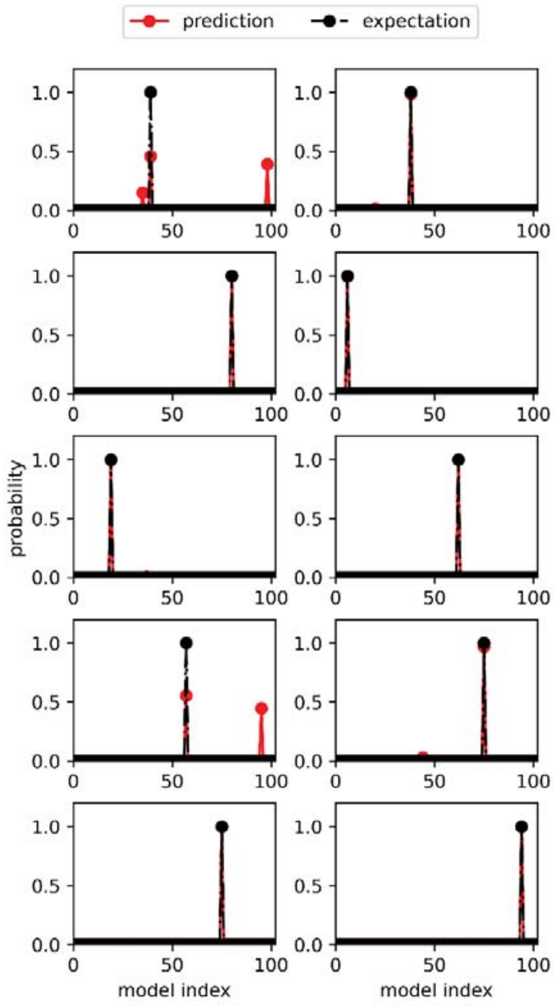

**Fig. SI 3.** Comparison of predicted (red lines) and expected (black dashed lines) model indexes for 10 different data sets. DLRN can well extrapolate the dynamics of systems, as also shown by the accuracy values in Table 1.

Fig. SI 4 shows a comparison of DLRN-predicted (red lines) and expected (black dashed lines) sorted time constant values for 10 different images. The x axis gives the index of each $\tau$ (index 0 = $\tau_1$, index 1 = $\tau_2$), while the y axis represents the corresponding time constants. The x axis has seven indexes (from 0 to 6) because each kinetic model can have a maximum of seven time constants due to the assumptions mentioned above. The plots in Fig. 3 demonstrate that DLRN predictions agree well with the expected curves, implying that DLRN can predict both the correct number of time constants and their corresponding values.

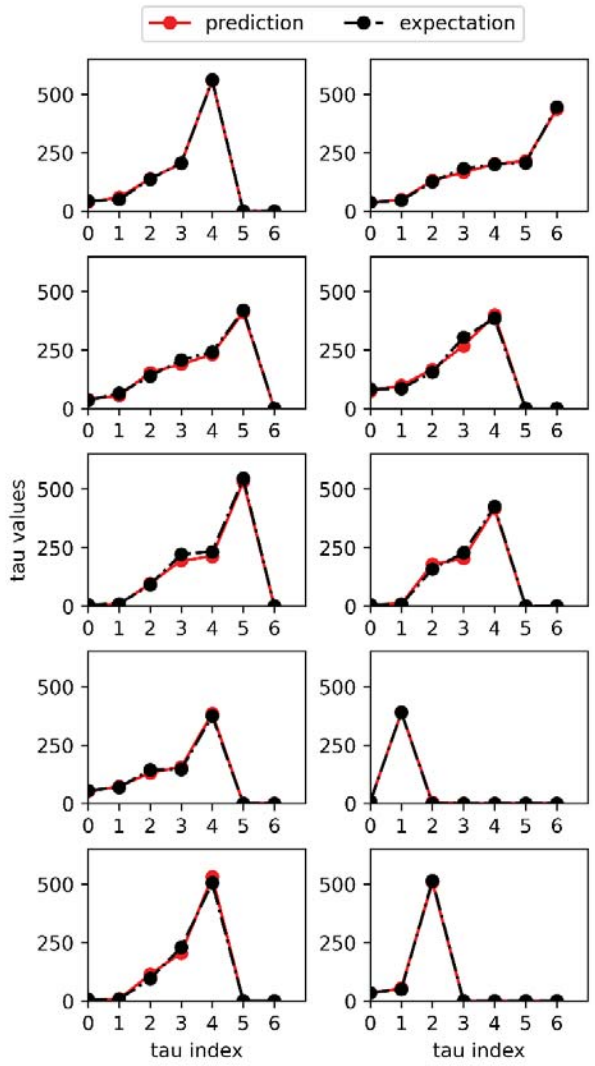

**Fig. SI 4.** Comparison of predicted (red lines) and expected (black dashed lines) time constants of specific pathways for 10 different data sets. DLRN analysis can extrapolate with good precision the number of time constants and their values. The accuracy of the DLRN, obtained using the area matrix, is shown in Table 1.

Fig. SI 5 shows a comparison between DLRN-predicted (continuous lines) and expected (dashed lines) spectral amplitudes for five different 2D datasets. In each graph, the x axis represents the wavelength, and the y axis shows the normalized

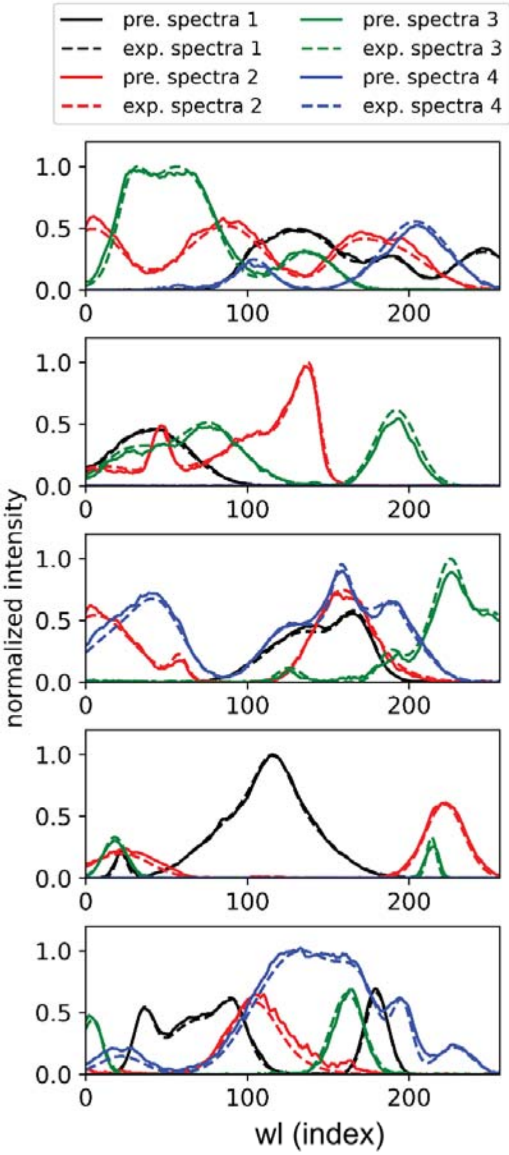


**Fig. SI 5.** Comparison of predicted (continuous lines) and expected (dashed lines) spectra of specific species for five data sets. DLRN can extrapolate the number of spectra (SAS) and their shapes with good approximation. The accuracy of the DLRN analysis is shown in Table 1 using the area metrics.

intensity of the amplitudes (normalized by the image's maximum). The maximum number of DLRN amplitude predictions is four due to the method of generating the data (see “Generation of synthetic data for training and evaluation” in the Methods section). DLRN also predicted the correct spectra for complex structures with good approximation.

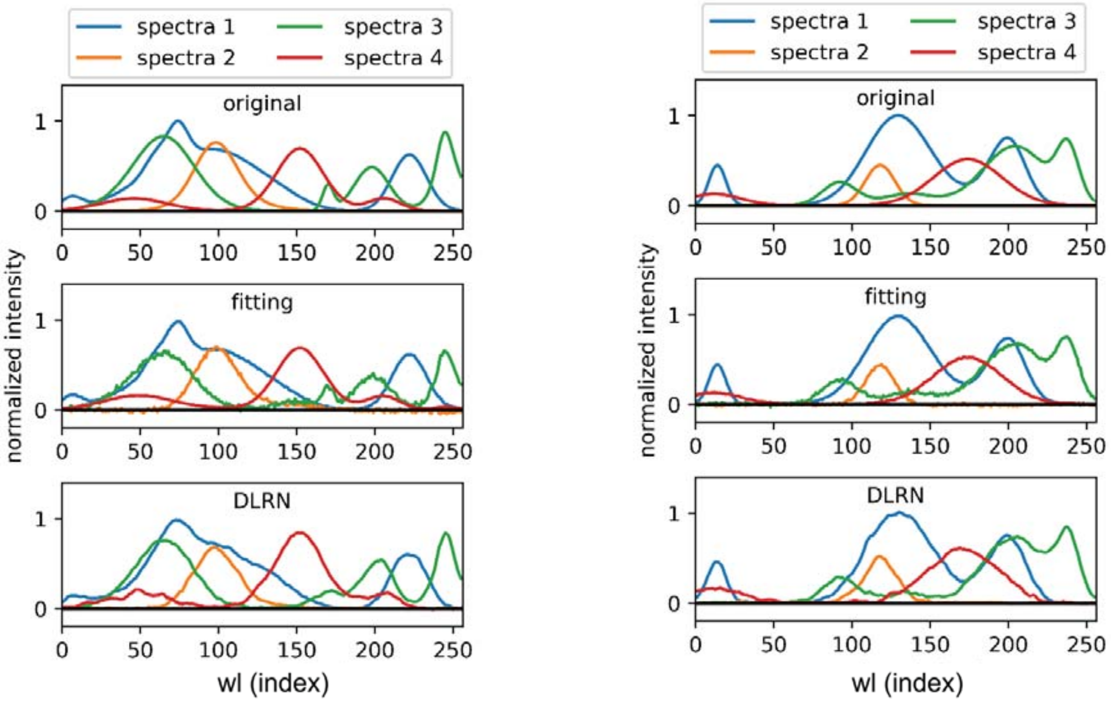


**Fig. SI 6.** Comparison of original spectra and the prediction from classical and DLRN analyses for two measurements. Classic and DLRN analyses show similar results, indicating that DLRN can perform spectral analysis well.

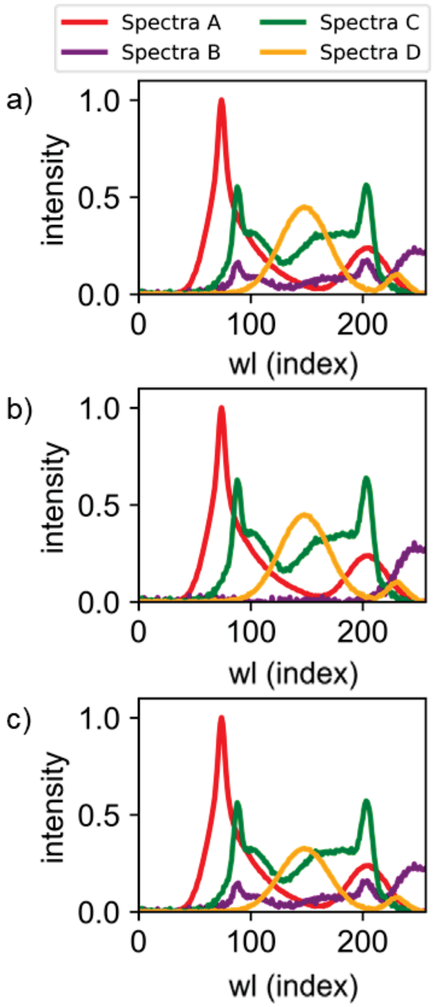


**Fig. SI 7.** Species-associated amplitude for three different fittings. The free solution shows the same residual values, but spectra and relative time constants differ depending on the initial fitting parameters. This makes multiple equal solutions mathematically possible, indicating the difficulty of correct model selection using classical analysis.

For a comparison, Figs. SI 5 and SI 6 show the expected spectra, the spectra obtained by fitting the data, and the DLRN analysis. In the main text, we show that DLRN sometimes performs better in cases of complex dynamics. Classical and DLRN analysis present very similar results for systems with more pronounced spectra, as shown in Fig. SI 6. Dependence of the classical results on the initial fitting parameters are shown in Fig. SI 7. These results support the efficiency and utility of DLRN in kinetic reaction networks analysis.

### 3. Generation of Training input and output datasets

We created a kinetic model to generate a synthetic 2D data set as an input for the training. Kinetic models can be expressed by a set of differential equations. For example, for the linear model A→B→GS (where GS is the ground state), we are considering only four possible active states, so the system of differential equations can be written in the following matrix form:

$$\begin{bmatrix} dA/dt \\ dB/dt \\ dC/dt \\ dD/dt \end{bmatrix} = \begin{bmatrix} -k_1 & 0 & 0 & 0 \\ k_1 & -k_2 & 0 & 0 \\ 0 & 0 & 0 & 0 \\ 0 & 0 & 0 & 0 \end{bmatrix} \begin{bmatrix} A \\ B \\ C \\ D \end{bmatrix} \quad \text{(Eq. SI 1)}$$

A, B, C, and D are the active states and $k_1$ and $k_2$ are the kinetic rate constants (which are the inverse of the time constants involved in the mechanism). The system of differential equations can be resolved by introducing the initial conditions, typically [A($t_0$),0,0,0], and assuming that initially only the active state A is populated. The solution gives us the concentration trace matrix **C**(t), which can be written like a system of equations:

$$\begin{cases} A(t) = A_0 e^{-k_1 t} \\ B(t) = A_0 \frac{k_1 - k_2}{k_1 k_2} \left(e^{-k_2 t} - e^{-k_1 t}\right) \\ C(t) = 0 \\ D(t) = 0 \end{cases} \quad \text{(Eq. SI 2)}$$

To complete the generation of the concentration trace matrix **C**(t), it is necessary to generate the values of the rate constants to be substituted within the equations. To this end, a set of about 210 time constants between 1 and 605 was generated with a nonconstant step (step = 1 between 1 and 20; step = 2 between 20 and 250; step = 5 between 250 and 600). Depending on the number of active states that contribute to generating the signal, variable subsets were created for each state. In our example of the linear mechanism above (Eq. SI 2), the only states contributing to the signal are A(t) and B(t), meaning that two subsets have emerged. This means that in the case of one active state, only one subset is generated (which in this case is the entire set); in the case of two active states, two subsets are created, and so on.

Fig. SI 8 shows how subsets are made variably. Starting with the creation of the subset for the state A(t), a random index is chosen between minimum and maximum values. After the selection, the subset for A(t) is made, and a new initial set is generated for the creation of the subsets for B(t), and this process is repeated for all the active states. The minimum and maximum values for index selection for A(t) were 25 and 82, respectively, whereas for the other states they are 30 and half of the length of the new initial set. The decay time constants were then selected from the appropriate subset for each active state. The reasons we decided to generate the tau values in this way are twofold: 1) we wanted each state to decay more slowly; 2) we wanted to generate as many cases as possible that satisfy condition 1. In the case of a parallel mechanism for an active state, the two decay time constants were also selected from the corresponding subsets, but with the condition that the quantum yield for a pathway is between 30% and 70%.

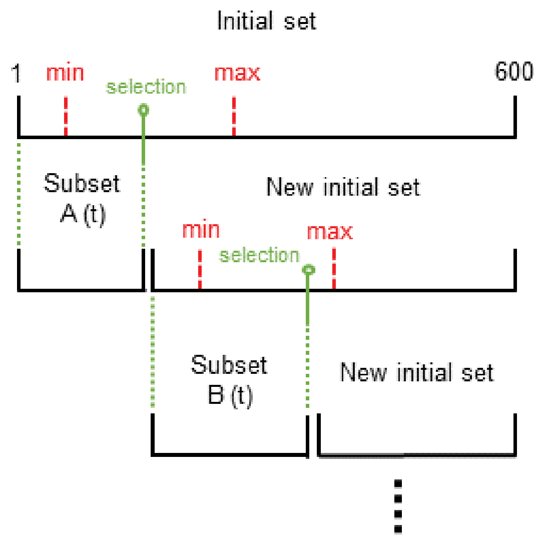


**Fig. SI 8.** Scheme for generating the dynamics subset for the active electronic state.

As mentioned in the main text (see Methods, Generation of synthetic data for training and evaluation), to generate the signal **X**(t,λ), we also need the amplitude matrix **S**(λ). For each active state, we needed to generate the corresponding spectrum (the amplitudes of the active state). To do so, each spectrum was generated with a sum of a random number of Gaussians between one and eight, where each Gaussian can have an amplitude between 0.2 and 1 and a width between 5 and 50. Notably, at least one Gaussian has a minimum amplitude of 0.4. After that, all spectra were collected to generate the amplitude matrix **S**(λ), where the first row is the amplitude vector of the active state A, the second row for the active state B, and so on.

The output training data were generated differently depending on the output block (model, time, and amplitude; Fig. SI 2). Starting with the model block, each kinetic model can be represented by a reference matrix **M**$_r$(*k*), which is linked to the *k* matrix part shown in Eq. SI 1. In detail, using the same k matrix as in Eq. SI 1, it is possible to write the **M**$_r$(k) for this kinetic model as follows:

$$\boldsymbol{M}_r(k) = \begin{bmatrix} -1 & 0 & 0 & 0 \\ 1 & -1 & 0 & 0 \\ 0 & 0 & 0 & 0 \\ 0 & 0 & 0 & 0 \end{bmatrix} \approx \begin{bmatrix} -k_1 & 0 & 0 & 0 \\ k_1 & -k_2 & 0 & 0 \\ 0 & 0 & 0 & 0 \\ 0 & 0 & 0 & 0 \end{bmatrix} \qquad \text{(Eq. SI 3)}$$

Here, the rate constants are replaced by the value 1, which indicates whether a state decays or is formed during the process. Another case to consider is that of a branching mechanism. Eq. SI 4 shows an example of how **M**$_r$(*k*) can be written in the case of a parallel mechanism:

$$\boldsymbol{M}_r(k) = \begin{bmatrix} -2 & 0 & 0 & 0 \\ 1 & -1 & 0 & 0 \\ 1 & 0 & 0 & 0 \\ 0 & 0 & 0 & 0 \end{bmatrix} \approx \begin{bmatrix} -(k_1 + k_2) & 0 & 0 & 0 \\ k_1 & -k_3 & 0 & 0 \\ k_2 & 0 & 0 & 0 \\ 0 & 0 & 0 & 0 \end{bmatrix} \qquad \text{(Eq. SI 4)}$$

Having the reference matrix for each of the 102 possible models we can generate, each **M**$_r$(*k*) is converted into a one-hot encoding vector and used as output training for the neural network.

In the case of the time block, it is necessary to introduce a second input for the analysis and generate a correct output data set. For this purpose, we used the **M**$_r$(*k*) matrix (see Fig. SI 2). A vector of seven values was generated using the output label, in which each element is the time constant value of each decay pathway. Using the kinetic model shown in Eq. SI 4, we obtained the tau vector in a specific order:

$$model = \begin{bmatrix} -(k_1 + k_2) & 0 & 0 & 0 \\ k_1 & -k_3 & 0 & 0 \\ k_2 & 0 & 0 & 0 \\ 0 & 0 & 0 & 0 \end{bmatrix} k_1 < k_2 \qquad \text{(Eq. SI 5)}$$

$$out_{tau} = \left[\frac{1}{k_1}, \frac{1}{k_2}, \frac{1}{k_3}, 0,0,0,0\right] \qquad \text{(Eq. SI 6)}$$

The amplitude block works similarly to the time block, which uses the **M**$_r$(k) matrix as a second input (see Fig. SI 2). However, the output has a 256 × 4 matrix in which each row is the spectrum of the corresponding electronic state (first row for A(t), second row for B(t), and so on).

### 4. Selection of the area metrics limits

The area metric $A_M$ was introduced to evaluate the accuracy of DLRN in predicting the values of the time constant and amplitude of a kinetic model. This metric requires a threshold value to determine whether a measurement can be considered positive or not (Methods, Area metrics for regression accuracy). For this study, we chose values of 0.8 and 0.9 for evaluating the time constant and a value of 0.8 for the amplitude prediction. The choice of these values can be explained using Fig. SI 9 as an example. The top graph shows a set of expected tau values (black line) and the area of values (blue area) within ± 20% error of the expected values. This area corresponds to all the possible values a DLRN prediction can have and yield a value of $A_M$ greater than 0.8. By holding this value as a threshold for positive values, we consider all predictions to have an error on a single tau of less than 20%.

In the case of amplitude calculation, four spectra are predicted using DLRN. This means that the 20% error obtained using $A_M > 0.8$ must be divided for each spectrum, indicating an average error of 5%.

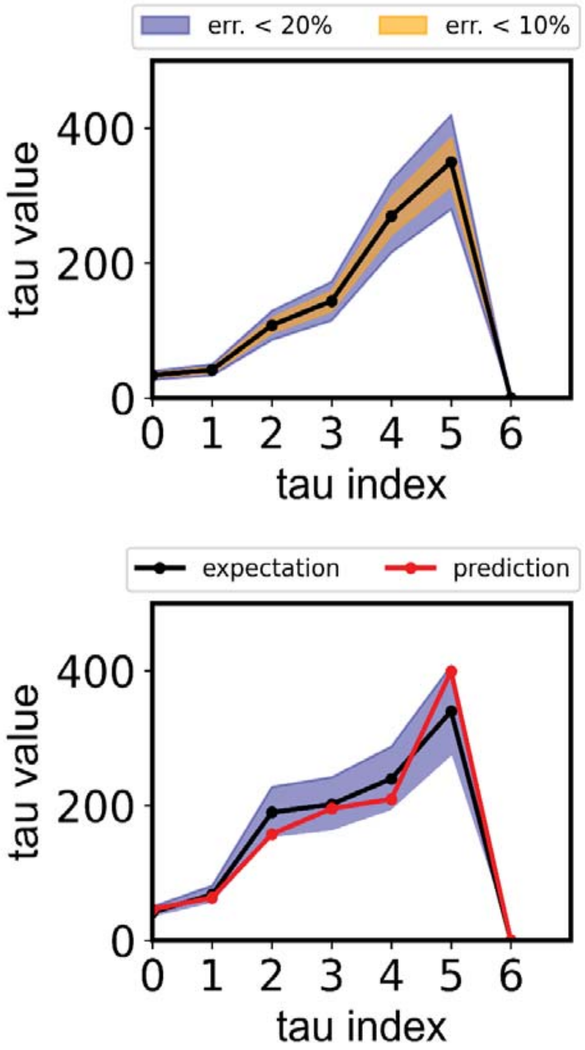


**Fig. SI 9.** Top: Set of time constants expectation value (black line). The blue area shows all the values within an error of 20% compared to the expectation. Bottom: Example of a DLRN prediction (red line) having $A_m$ = 0.85 in comparison to the expectation. The blue area shows all the values within an error of 20% compared to the expectation.

### 5. DLRN Analysis on time-resolved agarose gel

This section shows the model, time constants, and amplitude prediction on time-resolved agarose gel, as shown in the main text for the time-resolved spectra.

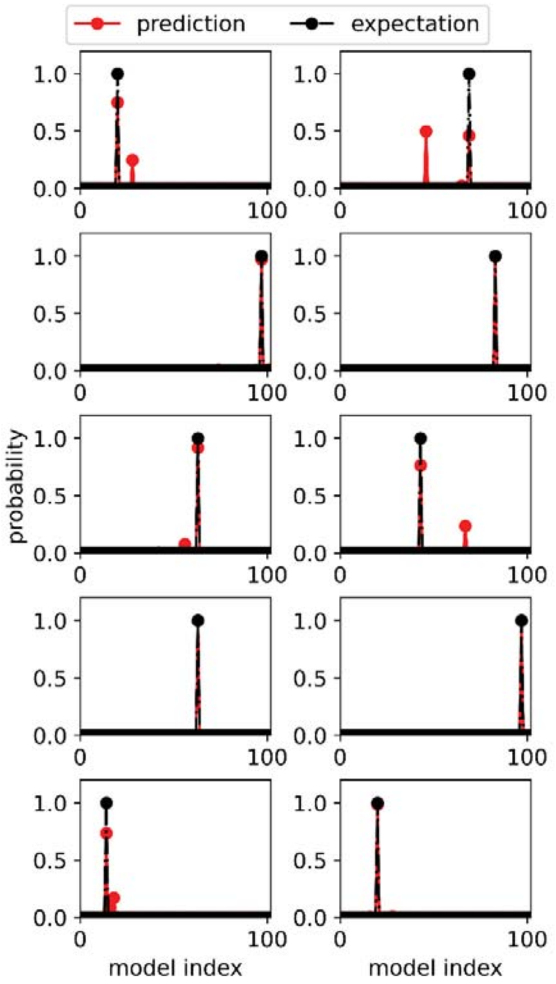


**Fig. SI 10.** Comparison of predicted (red lines) and expected (black dashed lines) model indexes for 10 different agarose gel images. DLRN can extrapolate the dynamics of systems well, as the accuracy values in Table 2 also show.

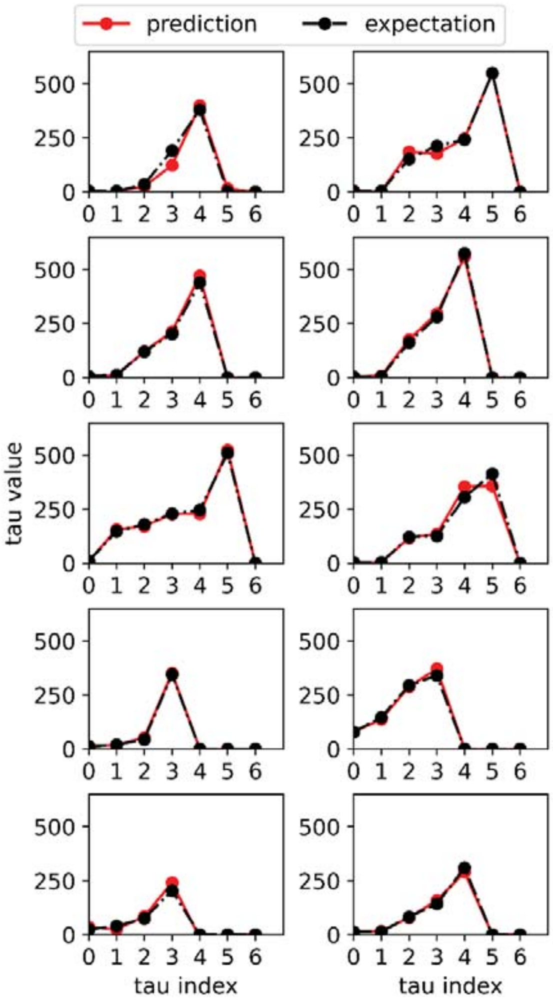


**Fig. SI 11.** Comparison of predicted (red lines) and expected (black dashed lines) time constant values of specific pathways for 10 different agarose gel images. DLRN can extrapolate with good precision the number of time constants and their value during the analysis. The accuracy of DLRN is shown in Table 2 using the area metric.

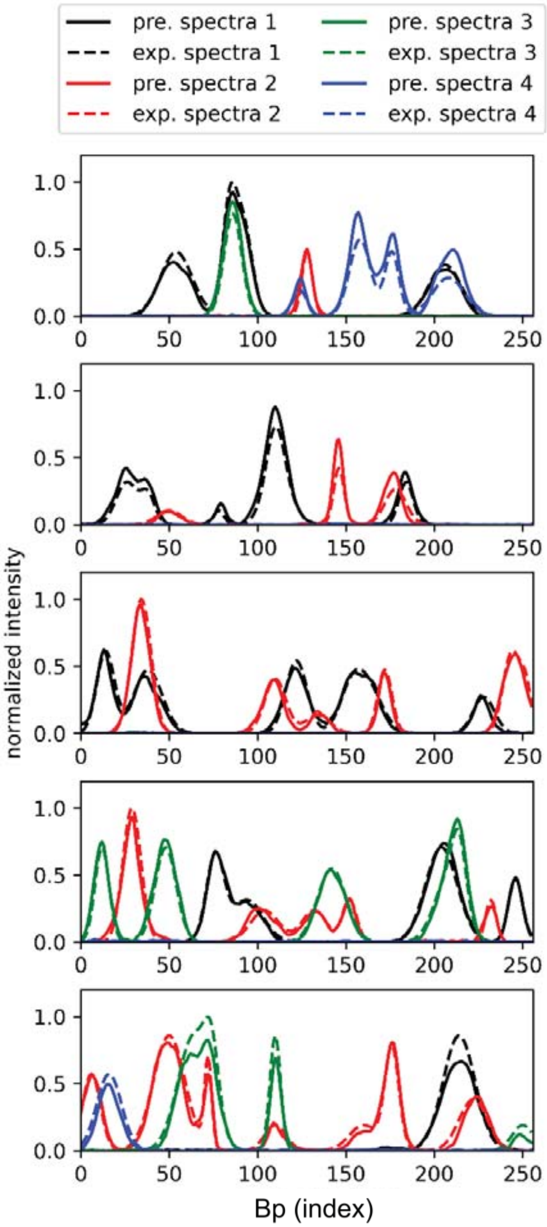


**Fig. SI 12.** Comparison of predicted (continuous lines) and expected (dashed lines) spectra of specific species for five agarose gel images. Each chart shows one data set. DLRN can extrapolate with good approximation the number of time spectra and their shapes. The accuracy of the DLRN analysis is shown in Table 2 using the area metrics.

### 6. DNA strand displacement: additional analysis

Fig. SI 13 shows the expected and forced kinetic traces of the two DSD kinetic reactions shown in Fig. 6 of the main text. Forced kinetic traces were obtained by using a combination of DLRN predictions (for $\tau_{A\to B}$ and $\tau_{A\to C}$) and calculated (for $\tau_{C\to D}$ and $\tau_{B\to D}$) tau values. The graph shows that expected and forced kinetic traces diverge significantly when the calculated $\tau_{C\to D}$ and $\tau_{B\to D}$ are used, pointing that DLRN analysis was correct.

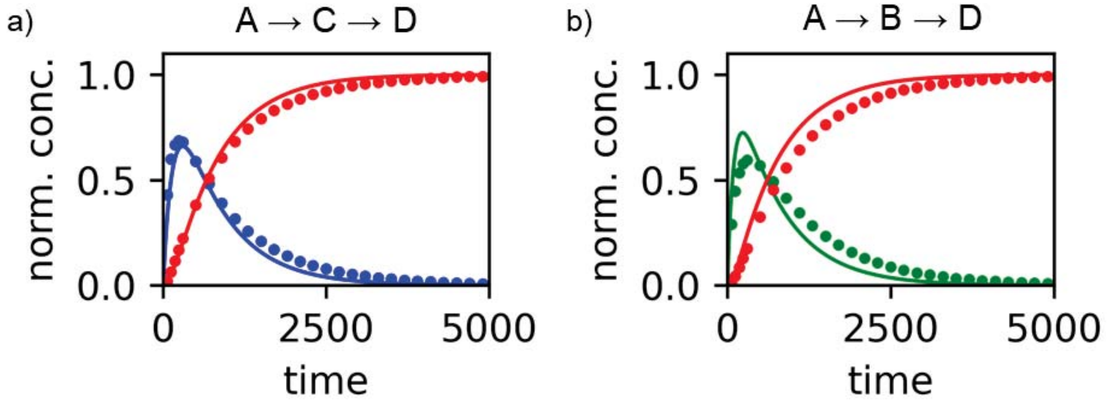


**Fig. SI 13.** Expected (circles) and forced (continuous lines) kinetic traces for the state C (blue), B (green), and D (red) for DSD kinetic reaction A→C→D (a) and A→B→D (b). The forced kinetic traces are a combination of DLRN predictions for the initial reaction (A→C and A→B), whereas for the second reaction step (C→D and B→D) the expected value of 721 s was used. The predicted kinetic traces differ significantly from the expected values, suggesting that DLRN tau predictions were correct.

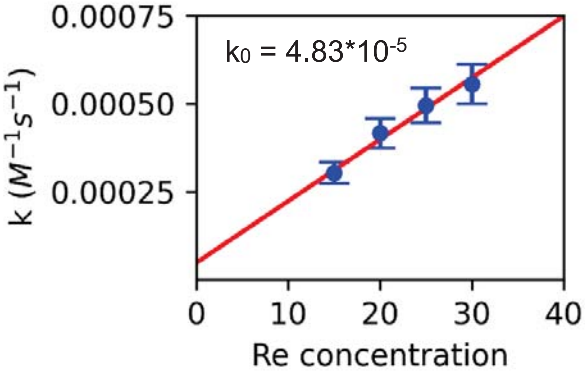


**Fig. SI 14.** Rate constant $k_{C\to D}$ predicted with DLRN for different concentration values of Re. Imposing an error of 10% (DLRN maximum error), the linear fit gives an intercept value ($k_0$) of $4.83 \times 10^{-5}$, which is similar of the expected rate constant of $4.62 \times 10^{-5}$. The difference in these two values is approximately 4.5%.

Fig. SI 14 shows the rate constants for C→D kinetic pathways obtained from DLRN analysis using four Re concentrations and then fitted by a linear regression. The intercept represents the value with no Re present, which can be compared to the theoretical value of $k_0 = 4.83 \times 10^{-5}$. The linear regression shows an intercept value of $4.62 \times 10^{-5}$, which is only 4.5% different than $k_0$. Fig. SI 15a shows the residuals obtained by DLRN analysis of the full DSD dynamics if both Su1 and Su2 are present. The errors of the residuals were not significant, indicating the predictions from the DLRN were good. Fig. SI 15b presents the normalized kinetic traces for states B and C, showing the remarkably similar dynamics of the two states and demonstrating that disentangling the two contributions is extremely difficult.

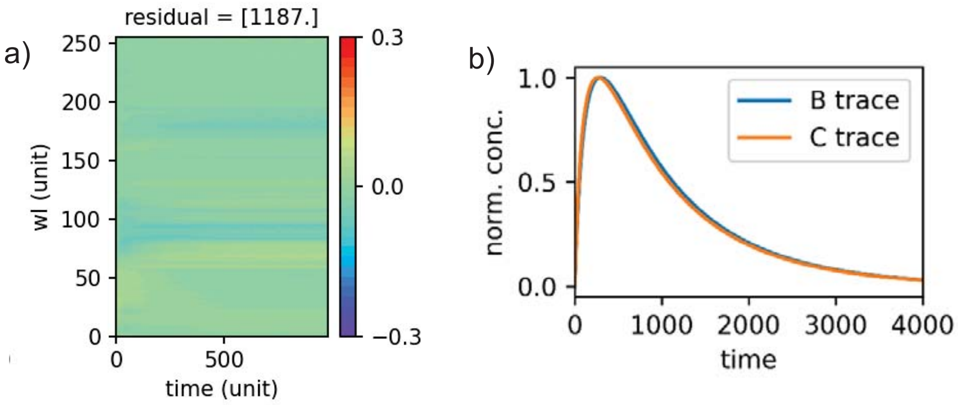


**Fig. SI 15. a,** Residuals obtained from DLRN analysis. **b,** Expected kinetic traces for the states B and C (see Fig. 7 in the main text) normalized by their respective maximum values. The two traces decay with the same time constants, making it difficult to disentangle the dynamics and ascertain the correct amplitude for the two states.

Fig. SI 16 shows the results of classical fitting on the DSD full dynamics. To fit the data, the following set of differential equations was used as a model:

$$\begin{bmatrix} dA/dt \\ dB/dt \\ dC/dt \\ dD/dt \end{bmatrix} = \begin{bmatrix} -k_1 & 0 & 0 & 0 \\ \alpha * k_1 & -k_2 & 0 & 0 \\ (1-\alpha) * k_1 & 0 & -k_3 & 0 \\ 0 & k_2 & k_3 & 0 \end{bmatrix} \begin{bmatrix} A \\ B \\ C \\ D \end{bmatrix} \qquad \text{(Eq. SI 7)}$$

where alpha is a fitting coefficient that can vary between 0.01 and 1. Here, fitting gave an alpha value of 0.26. Despite the use of the correct hypothetical model for the fit, the classical fitting yielded worse results than DLRN, albeit with slightly cleaner amplitude values.

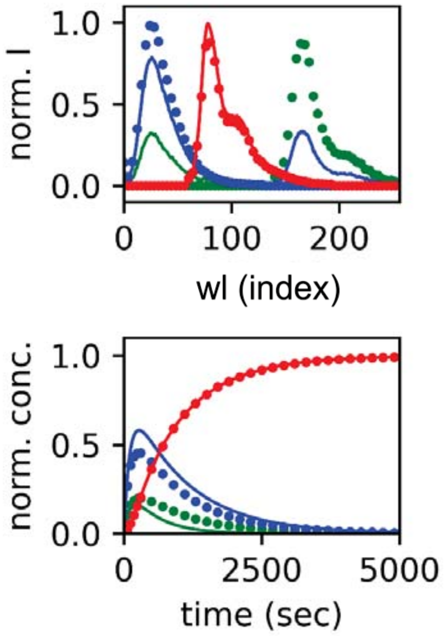


**Fig. SI 16.** Expected (circles) and classically fitted (continuous lines) amplitudes (top) and kinetic traces (bottom) for the branching kinetic reaction (Fig. 7 in the main text). The colors correspond to the states in Fig. 7. To fit the data, a branching mechanism and only positive amplitudes were imposed. Surprisingly, with respect to the kinetic traces the fitting performed worse than DLRN, but a cleaner amplitude prediction was obtained. However, the expected and predicted spectra for state B (green) significantly differ.

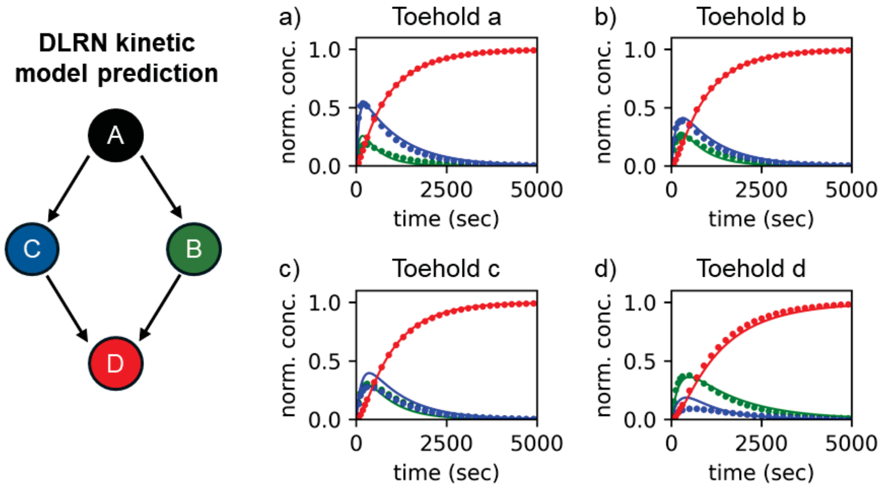


**Fig. SI 17.** Left: DLRN kinetic model prediction obtained by analyzing the four CRN variants having toeholds a–d. Right: Expected (circles) and DLRN-predicted (continuous lines) kinetic traces if toeholds a, b, c, and d (a–d, respectively) were used. Colors correspond to the kinetic model.

### 7. G-User-DLRN: a graphic user interface for DLRN analysis

This section provides a protocol for using the DLRN GUI for data analysis. The steps for running the GUI after starting the program called "DLRN_GUI" are given below. A new window appears from which a number of options can be selected (Fig. SI 18a).

1. Select Spectra or Agarose Gel from the first set of options. This will load the pretrained DLRN model for the respective scenario. This can take a few minutes.
2. Select Top 1 or Top 3 from the second set of options. This will change the analysis output, giving the solution for either the most probable output or the three most probable outputs.
3. Select the scale factor (suggested value = 1). This rescales the timescale to let DLRN analyze data sets with a time window larger than one timescale. However, using a large value for the scale factor can change the results of the analysis due to data interpolation during preprocessing data preparation.
4. Load the timescale to be used for the measurements. It is important to rescale the data with one that matches the timescale used during the DLRN analysis.
5. Load the data to be analyzed using "load the data" (located at the bottom). Search for the data that you want to analyze using the browser window. Only a NumPy zip file (.npz) having a subfolder "train" or .txt/.dat files can be loaded in the GUI.
6. (Optional) Is it possible to test the DLRN performance using the "Test DLRN" button. This allows the user to try a few ground truth data to check the performance.
7. Click the "Data Analysis" button to start the analysis and obtain the DLRN analysis results. This can be done only after the compulsory steps (1–5) have been completed. A typical output of the analysis is shown in Fig. SI 18b–d.

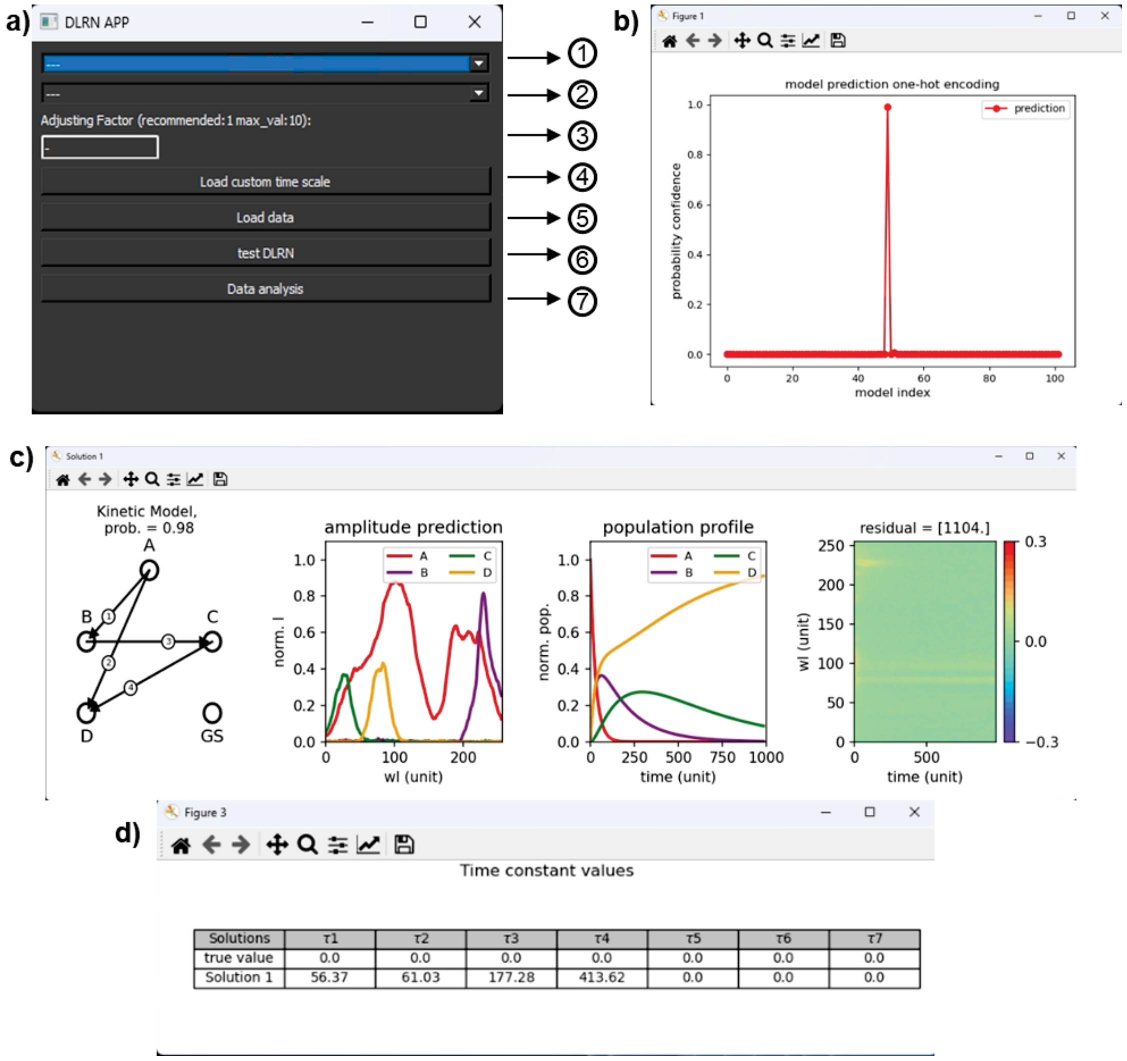


**Fig. SI 18. a,** The G-user-DLRN interface. The options 1–7 are outlined in the instructions provided above. **b–d,** The results obtained by DLRN analysis using the Top 1 solution: the one-hot encoding representation (b); the predicted kinetic model, tau prediction, and so forth (c); the table of tau values (d). If Top 3 analysis is selected, three windows like that shown in panel c appear, each detailing the DLRN results for their respective solution.

### 8. DNA sequences and corresponding reaction rates in the DSD

We calculated the binding rate coefficients for toeholds ($k_{\text{bind}}$) using Eq. SI 10, with equilibrium concentrations obtained from NUPACK, assuming $k_{\text{unbind}} = 0.012\ \text{s}^{-1}$, $[Na^+]$ = 100 mM, $[Mg^{2+}]$ = 10 mM, and 37 °C.

$$\text{t1} + \text{t1}^* \rightleftharpoons \text{t1t1}^* \qquad \text{(Eq. SI 8)}$$

$$K_{eq} = \frac{k_{\text{bind}}}{k_{\text{unbind}}} = \frac{[\text{t1t1}^*]}{[\text{t1}][\text{t1}^*]} \quad \text{(Eq. SI 9)}$$

$$k_{\text{bind}} = \frac{[\text{t1t1}^*]}{[\text{t1}][\text{t1}^*]} k_{\text{unbind}} \quad \text{(Eq. SI 10)}$$

However, this estimation does not take into account the significant effect of fluorophores and quenchers. The results of preliminary experiments have suggested that the real processes are much faster, therefore we used 100-fold higher values for $k_{\text{bind}}$. The values are summarized in Table SI 2.

**Table SI 1.** Realistic DNA sequences for the above DNA reaction network.

| Name | Structure | Sequence |
|---|---|---|
| Inp | 1 2 3 4 | a: AATAATCTACC TCAGCACATCGTATCAAACTCGTCCATGGGTTAGG<br>b: CTAATCTACCTCAGCACATCGTATCAAACTCGTCCATGGGTTAGG<br>c: GCTCTACCTCAGCACATCGTATCAAACTCGTCCATGGGTTAGG<br>d: AATCTACCTCAGCACATCGTATCAAACTCGTCCATGGGTTAGG |
| Su1 | 2 5 3 6<br>6*3*2*1* | TCAGCACATCCTGTTCGGTATCAAACTCGTCCAT\Atto425\TGTC<br>a: GACA\Q\ATGGACGAGTTTGATACGATGTGCTGAGGTAGATTATT<br>b: GACA\Q\ATGGACGAGTTTGATACGATGTGCTGAGGTAGATTAG<br>c: GACA\Q\ATGGACGAGTTTGATACGATGTGCTGAGGTAGAGC<br>d: GACA\Q\ATGGACGAGTTTGATACGATGTGCTGAGGTAGATT |
| Su2 | 7 2 5 3<br>8*4*3*2*7* | ATAGTCAGCACATCCTGTTCGGTATCAAACTCGTCCAT\Cy5.5\<br>CCTAACCC\Q\ATGGACGAGTTTGATACGATGTGCTGACTAT |
| Re | 3<br>3*5* | GTATCAAACTCGTCCAT\Cy3\<br>\Q\ATGGACGAGTTTGATACCGAACAG |

Q indicates a wide-range quencher. The spectra used in the analysis were generated according to the fluorophores used: Atto425, Cy5.5, and Cy3.

**Table SI 2.** Binding rate coefficients for the different toeholds.

| Toehold name | Sequence | $k_{\text{bind}}$ ($nM^{-1}s^{-1}$) |
|---|---|---|
| a | AATAATCTACC | $1.43 \times 10^{-3}$ |
| b | CTAATCTACC | $5.73 \times 10^{-4}$ |
| c | GCTCTACC | $2.94 \times 10^{-4}$ |
| d | AATCTACC | $5.58 \times 10^{-5}$ |
| 4 | GGGTTAGG | $3.29 \times 10^{-4}$ |
| 5 | CTGTTCG | $4.62 \times 10^{-5}$ |
